# Small Molecule Modulation of MHC-I Surface Expression: A Click Chemistry-Based Discovery Approach

**DOI:** 10.1101/2025.01.31.635109

**Authors:** Sarah E. Newkirk, Joey J. Kelly, Mahendra D. Chordia, Yue Dou, Tian Zhang, Marcos M. Pires

## Abstract

Immunotherapy has emerged as a powerful strategy for combating cancer by harnessing the patient’s immune system to recognize and eliminate malignant cells. Major histocompatibility complex class I (MHC-I) plays a pivotal role by presenting neoantigens to CD8+ T cells, triggering T cell-mediated killing. However, cancer cells often evade detection by downregulating MHC-I surface expression, hindering the immune response. This resistance mechanism offers an opportunity to bolster MHC-I surface expression *via* therapeutic interventions. We conducted a comprehensive evaluation of previously purported small molecule MHC-I inducers and identified heat shock protein 90 (Hsp90) inhibitors as privileged enhancers. Using a core scaffold, we employed an *in situ* click chemistry-based derivatization strategy to generate 380 novel compounds. New agents showed high induction levels, with one triazole-based analog, **CliMB-325**, also enhancing T cell activation and exhibiting lower toxicity. Altogether, we demonstrated the potential of click chemistry-based diversification for discovering small molecules to counter immune evasion.

## INTRODUCTION

The immune system can, often times, be remarkably precise, efficient, and powerful in detecting and eliminating cancerous cells. One of the principal mechanisms that the immune system leverages for cancer cell detection involves the presentation of cancer-specific peptides on the major histocompatibility complex (MHC) of the pathogenic cell.^1^ In particular, MHC class I (MHC-I) is a membrane protein expressed on most nucleated cells and is responsible for presenting short peptides (typically 8-12 amino acids in length) to CD8+ T cells.^2^ Recognition of a peptide-MHC complex (pMHC) by a CD8+ T cell *via* its T cell receptor (TCR) can result in a cytotoxic response through the release of a host of agents including perforin and granzyme B.^3^ For a CD8+ T cell to be activated against a target cell (and undergo subsequent phenotypic changes), it must first recognize a ‘non-self’ peptide which is generated inside the cell and presented on MHC-I. The non-self peptides that are typically found on the surface of cancer cells are broadly referred to as neoantigens.^4^

Neoantigens are generated *via* structural alterations to the proteome of cancer cells through amino acid substitutions,^5^ post-translational modifications,^6^ and other mechanisms.^7^ These non-self peptides, which can be loaded on MHC-I for presentation, may engage with TCRs on T cells in ways that are distinct from the unmodified peptides.^8^ Therefore, CD8+ T cells displaying the cognate TCR are well-positioned to specifically recognize and respond to neoantigen-presenting cancer cells, thereby promoting an anti-cancer immune response.^7^ In many instances, these mechanisms are central to eliminating the emergence of cancerous cells. Yet, there is considerable evidence demonstrating that cancer cells can actively avoid immune recognition by CD8+ T cells.^9, 10^ These mechanisms of resistance include, but are not limited to, remodeling of the tumor environment to create hypoxic and immunosuppressive conditions,^11–13^ increasing the expression of immune checkpoint proteins (e.g., programmed death-ligand 1 [PD-L1]),^14, 15^ promoting the secretion of immunosuppressive cytokines,^16^ and downregulating MHC-I molecules.^17^ Critically, the downregulation of surface MHC-I can impair patient responses to programmed cell death protein 1 (PD-1) blockade immunotherapy.^18^ Given the tremendous success of cancer immunotherapies that directly rely on MHC-I/TCR engagement, there is a clear need to discover potent agents that promote the expression of MHC-I in cancer patients. By restoring or enhancing MHC-I expression, it may be possible to overcome the immune resistance observed in many cancers, thus making cancer cells more susceptible to T cell-mediated killing (**Figure 1A**).

**Figure 1.**
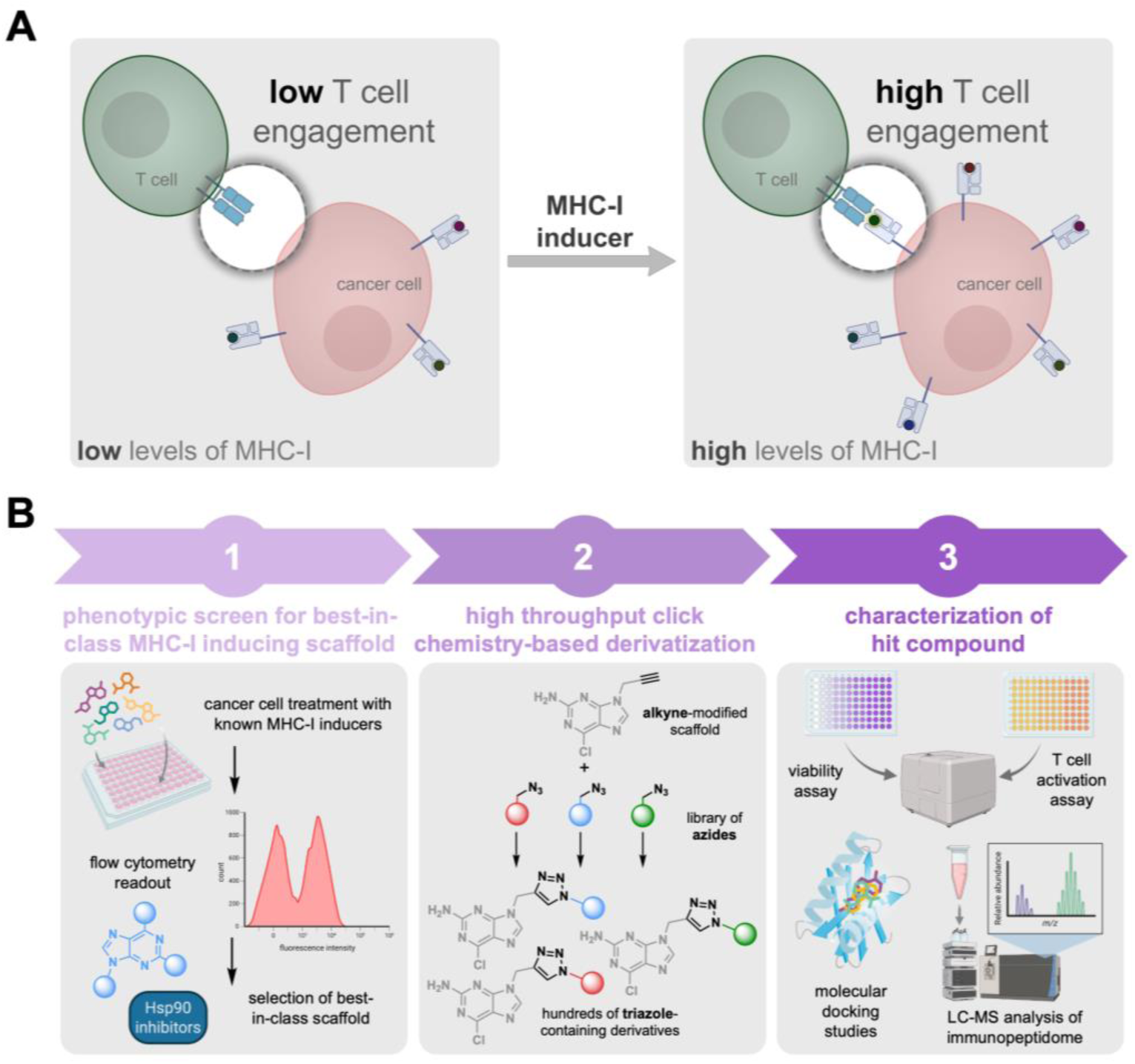
**(A)** Schematic representation of strategy to increase MHC-I expression and CD8+ T cell response. Cancer cells can downregulate the expression of MHC-I as a mechanism of evasion from a patient’s immunosurveillance. The use of small molecule inducers could potentially enhance immunotherapeutic approaches that are widely used in the clinic. **(B)** Workflow diagram illustrating the strategy and process used for the identification and discovery of small molecule MHC-I inducers.

The concept of using small molecules to promote the expression of MHC-I is not novel in and of itself, as a number of reports have identified compounds with this purported function. These compounds span a diverse range of biological activities and include DNA methyltransferase (DMNT) inhibitors,^19–26^ histone deacetylase (HDAC) inhibitors,^27–29^ kinase inhibitors,^30^ heat shock protein 90 (Hsp90) inhibitors,^31, 32^ stimulator of interferon genes (STING) agonists,^33–35^ and others.^36–39^ Although a wider range of FDA approved agents have been screened for MHC-I induction,^40, 41^ a definitive comparison across small molecule inducers has not, to the best of our knowledge, been previously reported. Given the diversity of reagents across the prior reports (e.g., cell lines, antibodies, concentrations, incubation times, etc.), it is critical to first establish the best-in-class scaffold. In this work, we conducted a rigorous head-to-head screen of small molecules to compare their ability to enhance MHC-I surface expression in colorectal cancer cells. Additionally, we demonstrate that Hsp90 inhibitors can increase the presentation of cancer-specific neoantigens. With a privileged scaffold in hand, we conducted a high-throughput diversification screen to generate a library of analogs that could be evaluated for their pharmacological properties **(Figure 1B**).

## RESULTS AND DISCUSSION

### Screening Small Molecules for their MHC-I Upregulation Activity

To identify the best-in-class drug scaffold for MHC-I upregulation, we utilized a flow cytometry-based assay. Briefly, CT26 murine colorectal cancer cells were incubated with individual compounds from a library of 25 small molecules that have previously been reported to enhance MHC-I expression. These included DNMT, kinase, HDAC, bromodomain and extra-terminal (BET), and Hsp90 inhibitors, as well as STING agonists and immunomodulatory drugs (IMiDs) (**Figure S1**). Following cellular treatment with each compound, a fluorescent anti-H-2K^d^ antibody was used to quantify MHC-I expression *via* flow cytometry (**Figure 2A**). As expected, most of these molecules demonstrated an increase in MHC-I surface expression at a high concentration of 5 μM (**Figure S2**). Still, it was notable that some of the molecules did not show any enhancement above background levels. To identify the most potent inducers of MHC-I surface expression, a second screen was conducted at a more stringent concentration of 1 μM. Our results revealed that three of the small molecules exhibited an increase in MHC-I surface expression above a two-fold cutoff (**Figure 2B**). These identified MHC-I enhancers fell primarily into two pharmacological classes: DNMT inhibitors (decitabine **1** and guadecitabine **3**) and Hsp90 inhibitors (zelavespib **23**). Further evaluation of the three lead compounds at a lower concentration of 500 nM showed that zelavespib (**23**) led to higher overall expression of MHC-I on the surface compared to the two DNMT inhibitors (**Figure 2C**). Based on these findings, we focused on Hsp90 inhibitors and expanded our search within this class beyond zelavespib in search of the most potent MHC-I inducers.

**Figure 2.**
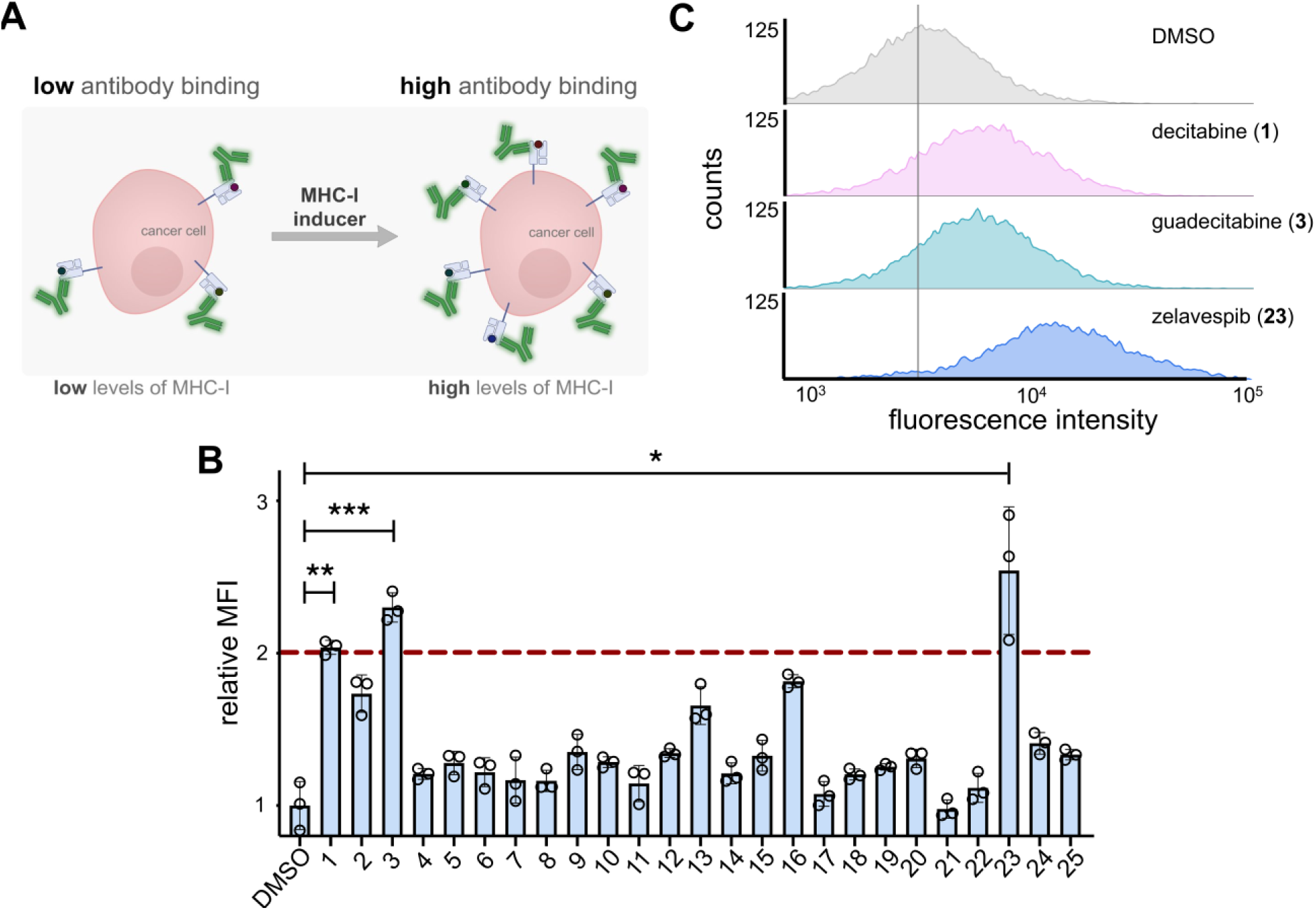
**(A)** Schematic representation of fluorescent antibody readout for increased MHC-I surface expression upon treatment with small molecules inducers. **(B)** Flow cytometry analysis of CT26 cells treated with 1 μM of indicated compounds (corresponding names and structures found in **Figure S1**). The red dashed line indicates threshold of 2-fold increase in MHC-I surface expression relative to DMSO control. MFI is the mean fluorescence intensity of the level of fluorescence relative to the DMSO control. Data are represented as mean ± SD (n=3). p-values were determined by a two-tailed *t*-test (* p < 0.05, ** p < 0.01, *** p < 0.001). **(C)** Flow cytometry histograms of CT26 cells incubated with 500 nM of indicated compounds. H-2K^d^ expression was measured by APC anti-mouse H-2K^d^ antibody. The vertical grey line represents median fluorescence intensity of DMSO treated cells. Data is shown as a representative histogram of the fluorescence intensity of 10,000 events per sample (n=3).

### Screening Hsp90 Inhibitors for MHC-I Surface Upregulation Activity

To further explore the relationship between Hsp90 inhibitors and MHC-I surface expression, six additional Hsp90 inhibitors were tested (for a total of seven Hsp90 inhibitors, including zelavespib from the primary screen of 25 small molecules) (**Figure 3A**). In total, this sub-library included three purine-based inhibitors, one resorcinol-based inhibitor, and three benzoquinone-based inhibitors. To more readily assess their potency, a concentration scan was performed in CT26 cells rather than a single-concentration analysis. Our results showed that all but two (pimitespib and tanespimycin) of the Hsp90 inhibitors tested had EC_50_ values in the nanomolar range for MHC-I surface expression. Among these, it was found that radicicol, BIIB021 and geldanamycin exhibited the lowest EC_50_ values, at 72 ± 1, 92 ± 1, and 144 ± 1 nM, respectively. Interestingly, these top hits represented all three primary classes of Hsp90 inhibitors, perhaps suggesting that Hsp90 inhibition is a primary driver of the phenotypic observation of MHC-I induction. Presumably, if any off-target activity were to be observed and if it were to be responsible for the MHC-I induction, it is likely that a single class would be favored over the others. We note that the top hits enhanced MHC-I surface expression in CT26 cells by approximately 5-to 8-fold relative to basal expression levels, representing a marked increase relative to the initial hit (zelavespib **23**) that prompted the focus on Hsp90 inhibitors (**Figure 3B**). To confirm the broader applicability of Hsp90 inhibitors as MHC-I inducers in other cellular contexts, we tested them in the human colorectal cancer cell line HCT116, where they also proved effective in enhancing MHC-I surface expression (**Figure 3C**). Overall, these results demonstrate that Hsp90 inhibitors are potent inducers of MHC-I surface expression. This prompted us to further explore Hsp90 inhibitors through the assembly of a larger and more diverse structure-activity relationship (SAR) campaign.

**Figure 3.**
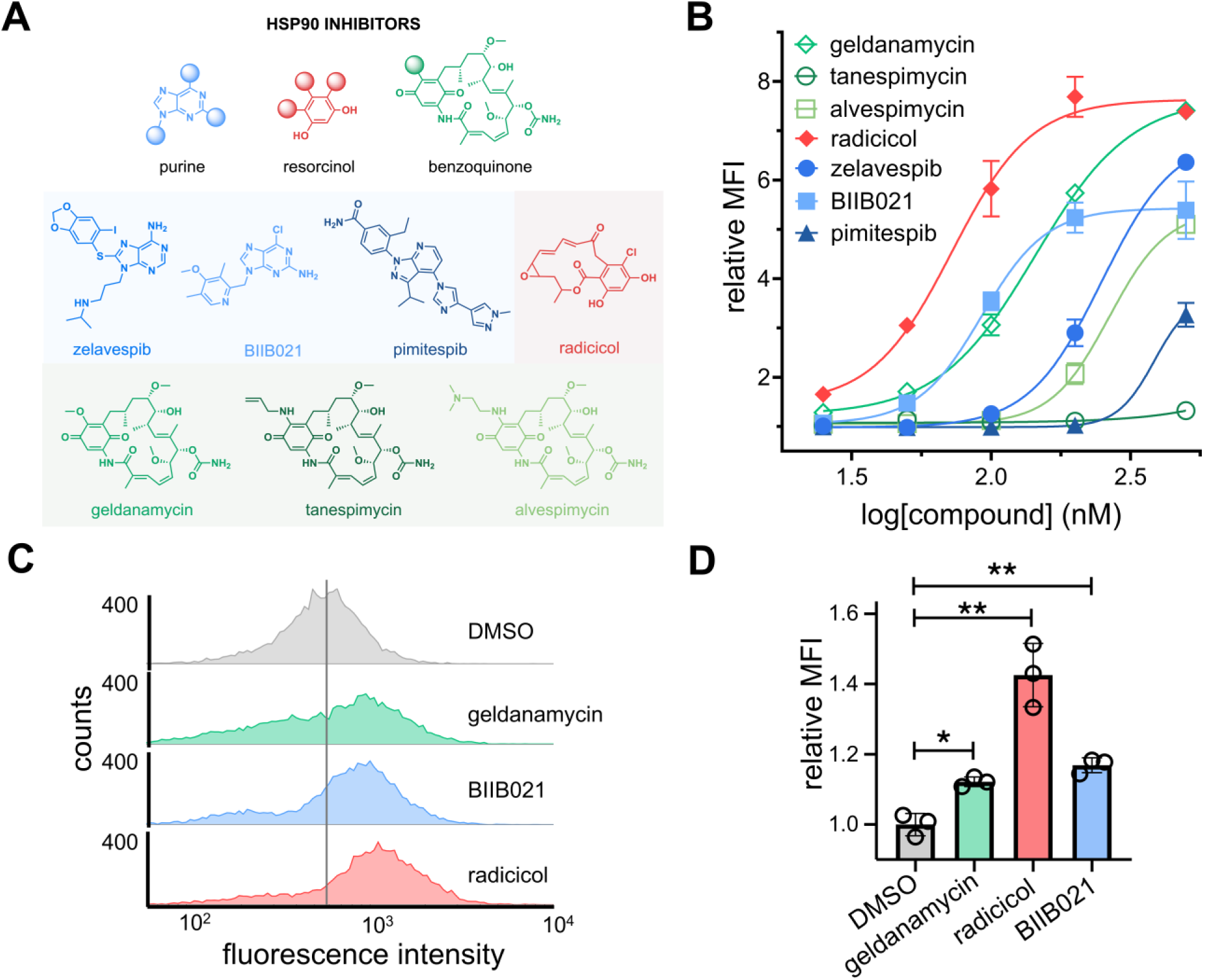
**(A)** Chemical structures of seven Hsp90 inhibitors tested for their enhancement of MHC-I surface expression. **(B)** Dose-response analysis by flow cytometry of CT26 cells treated with varying concentrations of seven Hsp90 inhibitors. H-2K^d^ expression was measured by APC anti-mouse H-2K^d^ antibody. Data are represented as mean ± SD (n=3), and Boltzmann sigmoidal curves were fitted to the data using GraphPad Prism. EC_50_ values are the concentration of compound needed to achieve 50% of the maximal MHC-I surface expression levels. **(C)** Flow cytometry histograms of HCT116 cells incubated with 200 nM of indicated compounds. HLA-A, B, C surface expression was measured by APC anti-human HLA-A, B, C antibody. The vertical grey line represents median fluorescence intensity of DMSO treated cells. Data is shown as a representative histogram of the fluorescence intensity of 10,000 events per sample (n=3). **(D)** Flow cytometry analysis of MC38-OVA cells treated with 100 nM of indicated compound. SIINFEKL-H-2K^b^ expression was measured by APC anti-mouse H-2K^b^ bound to SIINFEKL antibody. MFI is the mean fluorescence intensity of the level of fluorescence relative to the DMSO control. Data are represented as mean ± SD (n=3). p-values were determined by a two-tailed *t*-test (* p < 0.05, ** p < 0.01, **** p < 0.0001).

In theory, the enhancement of MHC-I surface expression should also enable the presentation of a greater breadth of cytosolic peptides, including potential neoantigens. This is particularly important because the efficacy of checkpoint blockage therapy depends primarily on cytotoxic CD8+ T cells recognizing neoantigens presented on the surface of cancer cells. As such, we next investigated the potential upregulation of specific antigens in live cells upon their treatment with Hsp90 inhibitors (**Figure 3D**). To test this, we utilized murine MC38-OVA cells, which are genetically modified to express the protein ovalbumin (OVA).^42^ OVA contains the sequence SIINFEKL, which is a well-established model neoantigen. Upon the intracellular processing of OVA resulting in the production of the SIINFEKL epitope, it is known that this peptide can be presented by H-2K^b^ and is recognized by SIINFEKL-specific CD8+ T cells.^43^ MC38-OVA cells were treated with 100 nM of the three lead Hsp90 inhibitors for 48 hours, followed by incubation with a fluorescent antibody specific for H-2K^b^ bound to SIINFEKL. Satisfyingly, treatment with Hsp90 inhibitors led to a significant increase in the presentation of the model neoantigen SIINFEKL, demonstrating that Hsp90 inhibitors can potentially promote the presentation of neoantigen-specific pMHC complexes (**Figure 3D**).

### High-Throughput Click Chemistry Diversification Strategy of Hsp90 Inhibitor

With the three top candidates in hand (radicicol, BIIB021, and geldanamycin), we sought to further diversify a core scaffold to broadly understand how structural features might drive MHC-I upregulation. We posed that developing a large-scale sub-library around a single agent could provide us with a wider set of compounds for testing and selection based on specific biological properties (e.g., improved toxicity profile, solubility, and selectivity). Among the top three Hsp90 inhibitors, we selected BIIB021 for further exploration due to its potency, the presence of structurally similar compounds in clinical evaluation for Hsp90 inhibition,^44–46^ and its robust chemical structure. Given the nature of our derivatization strategy, it was important to consider the potential stability of the starting scaffold. Both radicicol and geldanamycin contain structural fragments that are known to have low inherent chemical stability, making them less suitable for modification. Additionally, the availability of the crystal structure of BIIB021 in complex with Hsp90 can provide an avenue to understanding how analogs might interact with their target protein.^47^

For the library generation, we chose to use *in situ* click chemistry. In this format, an alkyne is installed within the core scaffold, and this alkyne-modified parent molecule is plated into a microwell plate system with each well containing a unique azide-tagged fragment. This approach offers considerable advantages for drug discovery of MHC-I inducers. Click reactions have a high level of specificity and efficiency, particularly exemplified by the Cu(I)-catalyzed azide-alkyne cycloaddition (CuAAC), which enables precise modifications and the synthesis of complex molecules with minimal byproducts.^48–50^ The inherent versatility in using a library of azides allows for extensive exploration of diverse molecular combinations and structural variations, which is essential for identifying drug candidates with optimal pharmacological properties. Moreover, this approach facilitates the simultaneous screening and synthesis of potential drug candidates, accelerating the identification of active compounds. Recently, this click chemistry-based strategy has been used to identify small molecule modulators of glucagon-like-peptide-1 receptor.^51^ Furthermore, it has been shown that over 80% of azide molecules formed triazole products with yields of 70% or higher using this method.^52^

In order to generate analogs of our core purine scaffold, we first needed to identify a site that was suitable for installing the alkyne handle. We appreciate that the change in MHC-I surface expression may be due to polypharmacological effects of Hsp90 inhibitors instead of a direct result of Hsp90 inhibition alone. However, since the increase in MHC-I surface expression was conserved across multiple classes of Hsp90 inhibitors, we used the crystal structure of BIIB021 complexed with Hsp90 to inform our decision on where to install the alkyne handle. Here, we identified the N9 position of the purine core as a solvent-exposed site that is amendable to chemical modification.^47^ Moreover, previous reports have demonstrated that this position can be leveraged to create purine-based analogs while preserving Hsp90 inhibition.^53^ Therefore, our envisioned approach involved modifying a precursor of BIIB021, 2-amino-6-chloropurine, by attaching an alkyne to the N9 position, enabling its reaction with a library of small molecule azides (**Figure 4A**). We hypothesized that the resulting triazole ring formed *via* the click reaction would structurally mimic the pyridine ring of BIIB021, allowing us to rapidly generate hundreds of derivatives for further exploration.

**Figure 4.**
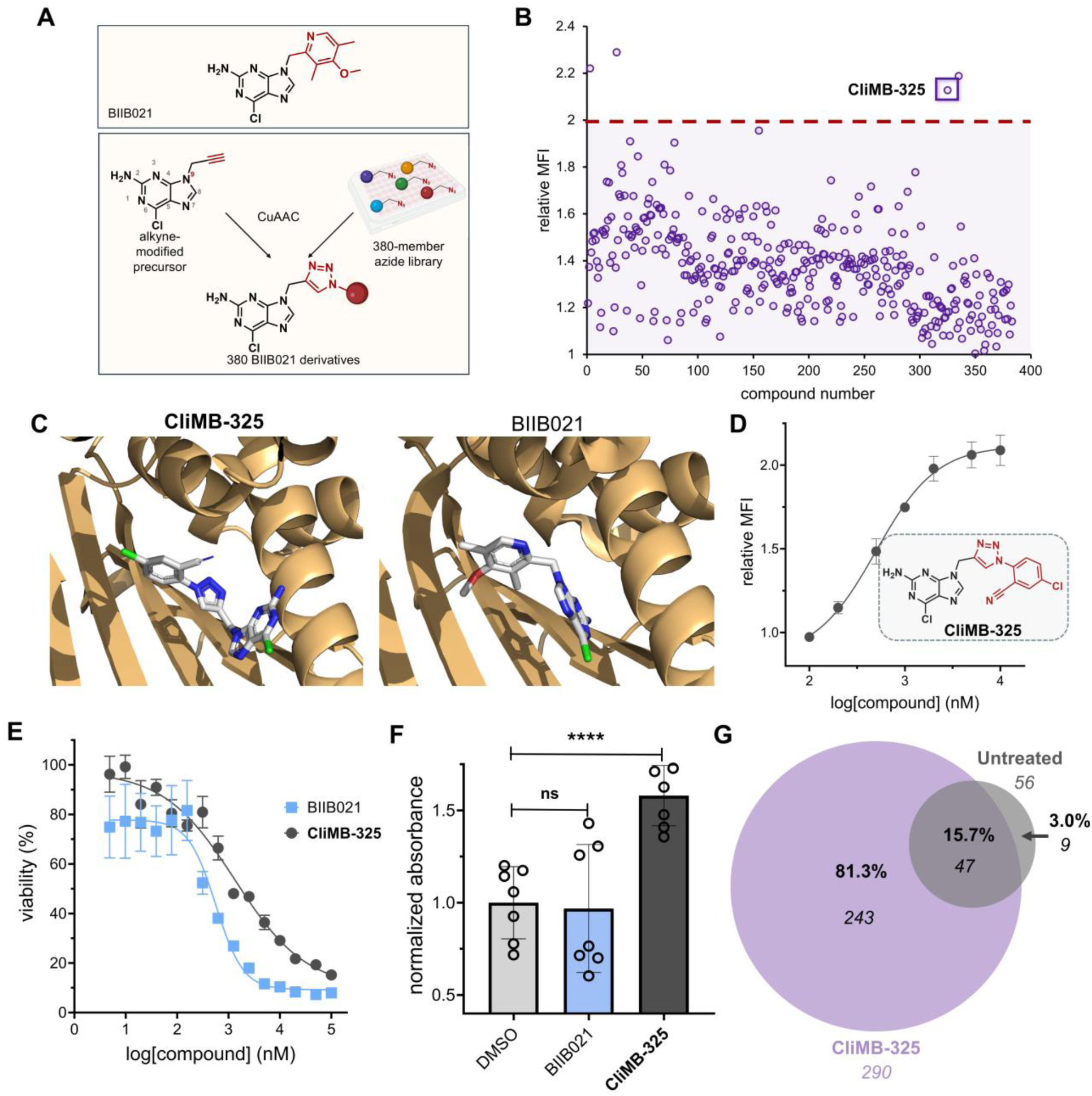
**(A)** Schematic representation of BIIB021 derivatization strategy. **(B)** Flow cytometry analysis of CT26 cells treated with click products between 9-propargyl-2-amino-6-chloropurine and the 380-member azide library. H-2K^d^ expression was measured by APC anti-mouse H-2K^d^ antibody and performed in singlet. Red dashed lined indicates threshold of 2-fold increase in MHC-I surface expression relative to DMSO control. MFI is the mean fluorescence intensity of the level of fluorescence relative to the DMSO control. **(C)** Stick models of **CliMB-325** (left) and BIIB021 (right) (white) docked into Hsp90 (light orange) were generated using existing crystal structure data (PDB ID: 3qdd) and Rosetta. **(D)** Dose-response curve and chemical structure of **CliMB-325**. CT26 cells were treated with varying concentrations of **CliMB-325**. H-2K^d^ expression was measured by APC anti-mouse H-2K^d^ antibody via flow cytometry. MFI is the mean fluorescence intensity of the level of fluorescence relative to the DMSO control. Data are represented as mean ± SD (n=3), and Boltzmann sigmoidal curves were fitted to the data using GraphPad Prism. EC_50_ values are the concentration of compound needed to achieve 50% of the maximal MHC-I surface expression levels. **(E)** Dose-response curves of CT26 cells treated with varying concentrations of **CliMB-325** or BIIB021 determined via MTT cell viability assay. Data are represented as mean ± SD (n=4), and nonlinear regression curves were fitted to the data using GraphPad Prism. CC_50_ values are the concentration of compound at which maximal cell viability is reduced by 50%. **(F)** MC38-OVA cells were incubated with 100 nM BIIB021 and 1 µM **CliMB-325** for 48 hours. Subsequently, cells were co-cultured with B3Z T cells for six hours. β-galactosidase expression was then measured via the colorimetric reagent CPRG on a plate reader at 570 nm. Data are represented as mean ± SD (n=7). p-values were determined by a two-tailed *t*-test (ns = not significant, **** p < 0.0001). **(G)** Venn diagram displaying MHC-I peptides eluted from CT26 cells *via* MAE that were exclusive to untreated (gray) or **CliMB-325**-treated (purple) cohorts. Peptides which were shared between groups are denoted in the overlapping region. Numbers in italics indicate total peptide numbers, while percentages in bold represent proportion out of 299 total peptides obtained. Data are derived from the same cell culture using 1 x 10^7^ CT26 cells.

To construct the alkyne-bearing purine, we reacted 2-amino-6-chloropurine with propargyl bromide to yield the major product, 9-propargyl-2-amino-6-chloropurine. The reaction also produced the N7 regioisomer as a minor product, which was separated during purification. The identity of isolated N9 alkyne-bearing compound was confirmed using NMR, and its purity was analyzed by RP-HPLC. Next, we sought to benchmark the reaction conditions to ensure their suitability for a larger screen. To do so, a model *in situ* CuAAC reaction was performed with the alkyne-modified precursor and a small subset of azide-containing molecules. In order to account for the structural variability likely to be found in the full library, we selected a subset of molecules that varied in size, polarity, and steric environment surrounding the azide group. The reactions were performed in microwell plates under experimental conditions designed to replicate those intended for the larger *in situ* reaction set. Notably, all three test reactions achieved a conversion rate of >90% to the triazole product (**Figure S3-S5**). With the model reactions showing high levels of conversion, we reasoned that the reaction conditions were well-suited for scaling up to a larger screen. The goal of utilizing a more extensive library was to ensure that modifications to the purine core would encompass a wide chemical space to broadly sample potential engagement with the target. Moreover, given the nature of the phenotypic assay, we reasoned that library diversity could also be important for improving other properties that are necessary for a lead candidate, such as high accumulation levels, low off-target effects, and reduced toxicity.

In total, 380 azide-containing small molecules (structures shown in **Figure S6**) were dispensed into wells containing the alkyne-bearing purine analog and the click reaction reagents. Following the reaction step, the contents of each well were incubated with CT26 cells, and MHC-I induction was assessed by treatment with a fluorescently tagged antibody as previously described. Critically, incubation of CT26 cells with the alkyne precursor alone or with the additional click reagents did not result in any increase in MHC-I surface expression, confirming that the reaction reagents themselves have no inherent activity (**Figure S8**). The results from the 380-member screen revealed that four of the click reaction mixtures (3, 27, 325, and 335), showed an increase in MHC-I surface expression above a 1.97-fold cutoff relative to the DMSO control **(Figure 4B)**. We selected this cutoff as it is a stringent three standard deviations above the mean response of all library compounds which would reduce the likelihood of obtaining a false positive hit. To further validate these initial results, cells were treated with each of the four reaction mixtures at a more stringent concentration (theoretically 500 nM, assuming complete conversion). Among these, treatment with compound 325 resulted in the highest levels of MHC-I surface expression (**Figure S9**). The click product formed between the alkyne precursor and compound 325 of the azide library was then synthesized and purified, yielding ‘**Cli**ck **M**HC-I **B**ooster-**325**’ (**CliMB-325**).

### CliMB-325 Enhances MHC-I Surface Expression, T Cell Activation, and Unique Peptide Display

To further characterize our novel purine-based lead agent **CliMB-325**, we utilized RosettaLigand to dock **CliMB-325** into Hsp90. The docking results revealed that **CliMB-325** binds to Hsp90 in a manner similar to BIIB021 (PDB ID: 3qdd) with no measurable deviation in the protein conformation (**Figure 4C**). Next, to establish its potency, a concentration scan of **CliMB-325** was conducted in CT26 cells to assess its ability to enhance MHC-I expression. Our findings demonstrated that **CliMB-325** retained MHC-I upregulation activity with an EC_50_ of 498 ± 1 nM (**Figure 4D**). Notably, the N7 position of the purine core in BIIB021 has also been shown as a site that tolerates chemical modifications while retaining Hsp90 inhibition activity.^45^ As such, we reacted azide 325 with the minor product of our alkyne-modified precursor (7-propargyl-2-amino-6-chloropurine) to synthesize the N7 isomer of **CliMB-325.** We found that this product exhibited comparable levels of MHC-I upregulation activity to **CliMB-325** in CT26 cells (**Figure S10**). Acknowledging the failed clinical progression of BIIB021,^44^ we also evaluated the viability of cells treated with **CliMB-325** compared to its parent compound, BIIB021, using an MTT cell viability assay. The results indicated that **CliMB-325** demonstrated a 2.4-fold improvement in toxicity over its parent compound, BIIB021, with CC_50_ values of 1.3 ± 0.0 nM µM and 563 ± 1 nM, respectively **(Figure 4E**).

Given that the downregulation of MHC-I surface expression impairs T cell recognition of neoantigens, we then sought to investigate whether the MHC-I upregulation induced by **CliMB-325** results in enhanced T cell activation. To do so, we utilized the B3Z T cell hybridoma cell line, which contains OVA-specific TCRs and expresses the enzyme β-galactosidase under the control of an IL-2 inducible promoter. Upon B3Z recognition of the OVA-pMHC complex on OVA expressing cells, the subsequent IL-2 production promotes the expression of β-galactosidase. β-galactosidase activity can then be measured *via* hydrolysis of the reagent chlorophenol red-β-galactopyranoside (CPRG) which leads to a color change that is reflective of T cell activation levels.^54^ MC38-OVA cells were treated with BIIB021 or **CliMB-325** for 48 hours and subsequently co-cultured with B3Z T cells. While BIIB021-treated cells showed no significant difference in TCR activation compared to the DMSO control, MC38-OVA cells treated with **CliMB-325** exhibited a nearly 1.6-fold increase in TCR activation levels (**Figure 4F**).

Building upon our finding that **CliMB-325** increases MHC-I surface expression in CT26 cells, we aimed to investigate potential changes to the immunopeptidome, including the enhancement of peptides presented on the cell surface. To this end, CT26 cells were incubated with or without **CliMB-325** for 48 hours, and MHC-I-bound peptides were isolated *via* mild acid elution (MAE) and sequenced using mass spectrometry.^55^ After removing known contaminants, the resulting peptide list was filtered to include only those with the optimal length (8-14 amino acids) for MHC-I binding, which we found constituted the majority of identified peptides (**Figure S11**).^56^ These peptides were further refined based on NetMHCpan 4.1 prediction scores to retain only putative binders of one or more of the three MHC allotypes expressed by the CT26 cells. Strikingly, **CliMB-325** treatment resulted in a 5-fold increase in the number of unique peptide sequences presented by MHC-I compared to untreated cells (**Figure 4G**). Among the 290 MHC-I peptides identified in the **CliMB-325**-treated sample, 81.3% were exclusive to this group and absent in untreated cells. Moreover, of the 56 total MHC-I peptides identified in the untreated group, 84% were also present in the **CliMB-325**-treated sample. This high degree of overlap in the immunopeptidome between the untreated and **CliMB-325**-treated samples suggests that **CliMB-325** treatment enhances the detection of peptides present in the existing immunopeptidome. Interestingly, the observed upregulation of surface MHC-I with **CliMB-325** treatment was modest (1.5-2-fold) compared to the impact on the immunopeptidome (5-fold), which may reflect a threshold effect in antigen presentation. While the concentration of peptides is generally the rate-limiting factor in the antigen presentation response, the relationship between MHC-I abundance and peptide diversity is nonlinear.^2^ Under conditions of limited MHC-I availability, peptide binding is highly competitive and high-affinity peptides dominate the repertoire. In this scenario, even a modest increase in available MHC-I can greatly (and disproportionately) expand the range of peptides successfully presented. Overall, these results validate the use of a high-throughput click chemistry screening approach to generate bioactive compounds with MHC-I upregulation activity.

## CONCLUSION

In this work, we have identified compounds with immunomodulatory effects on colorectal cancer cell lines. Among these molecules tested, Hsp90 inhibitors emerged as the most potent class of molecules for increasing MHC-I surface expression and promoting the display of cancer-specific neoantigens for CD8+ T cell recognition. Leveraging an Hsp90 inhibitor core scaffold, we have also demonstrated a proof-of-concept high-throughput click chemistry-based screening platform for the discovery of molecules with immunomodulatory activity. While our initial screen employed a 380-member azide library, we plan to expand to a 1,200-member azide library to sample a wider chemical space for molecules that upregulate MHC-I. Additionally, beyond the purine-based scaffold used in this study, we intend to extend this strategy to resorcinol- and benzoquinone-based scaffolds, which have also been favored for the development of new Hsp90 inhibitors.^57–62^ Ultimately, we envision that our approach of modifying preexisting chemical scaffolds with alkynes for large-scale click chemistry-based derivatization can be broadly applicable for various phenotypic screens beyond MHC-I upregulation, offering a versatile platform for drug discovery.

Counteracting cancer immune evasion mechanisms is essential to improving the efficacy of immunotherapy treatments. To this point, a substantial proportion of patients fail to respond to PD-1 blockade therapy due to the development of resistant tumors characterized by the downregulation of MHC-I.^18^ This challenge underscores the critical need for strategies that reengage the immune system by transforming immunologically ‘cold’ tumors back into ‘hot’ tumors that can be recognized and targeted for elimination. Therefore, we anticipate that the development of a widely applicable approach to enhance MHC-I surface expression represents a promising avenue to combat resistant cancers.

## EXPERIMENTAL SECTION

### Mammalian Cell Culture

CT26 cells were cultured in RPMI 1640 media supplemented with 10% fetal bovine serum, 50 IU/mL penicillin, and 50 µg/mL streptomycin. HCT116 cells were kindly provided by Dr. Anja-Katrin Bielinsky and were cultured in McCoy’s 5A media supplemented with 10% fetal bovine serum, 50 IU/mL penicillin, 50 µg/mL streptomycin, and 2 mM GlutaMAX. MC38-OVA cells were kindly provided by Dr. Mirna Perusina Lanfranca and were cultured in DMEM supplemented with 10% fetal bovine serum, 50 IU/mL penicillin, 50 µg/mL streptomycin, 50 µg/mL gentamycin, and 10 µg/mL blasticidin. B3Z cells were kindly provided by Dr. Aaron Esser-Kahn and maintained in RPMI 1640 media supplemented with 10% fetal bovine serum, 50 IU/mL penicillin, and 50 ug/mL streptomycin. All cells were cultured in T75 flasks and maintained in a humidified atmosphere of 5% CO_2_ at 37°C.

### Flow Cytometry-Based Assays

1.5 x 10^4^ cells were seeded in a treated 96-well plate along with indicated concentrations of library compounds at 37°C. After 48 hours, cells were washed once with PBS, removed using TrypLE^TM^ Express Enzyme (Thermo Fisher), and transferred to a round-bottom 96-well plate. Transferred cells were centrifuged (1100 x g, 5 min) in a Thermo Scientific Jouan C4i centrifuge, and the cell pellets were resuspended and fixed in 4% formaldehyde solution for 20 minutes. The plate was centrifuged (1100 x g, 5 min) and pelleted cells were resuspended in a 1:100 dilution of indicated fluorescence antibodies in culture media for 1 hour at 4°C. Flow cytometry was performed using the following antibodies: APC anti-mouse H-2K^d^/H-2D^d^ (clone 34-1-2S), APC anti-human HLA-A,B,C (clone W6/32), or APC anti-mouse H-2K^b^ bound to SIINFEKL (clone 25-D1.16). Cells were analyzed using an Attune NxT Flow Cytometer (Thermo Fisher) equipped with a 637 nm laser with 670/14 nm bandpass filter. Populations were gated and no less than 10,000 events per sample were recorded.

### MTT Cell Viability Assay

1.5 x 10^4^ CT26 cells were seeded in a treated 96-well plate, either with or without compounds (BIIB021 and **CliMB-325**) at indicated concentrations at 37°C. After 48 hours, a solution of MTT in PBS (filter sterilized through a 0.2-µM filter) was added to each well to achieve a final concentration of 0.45 mg/mL. After incubating at 37°C for 2 hours, cells were centrifuged (1100 x g, 5 min) in a Thermo Scientific Jouan C4i centrifuge and the supernatant was removed. 100 µL of DMSO was added to each well to dissolve the formation of formazan precipitate. The absorbance of the solution in each well was read at 570 nm using a BioTek Synergy H1 Microplate Reader. Wells containing no cells (only the added DMSO) were used as a negative control for viability, while untreated cells served as the positive control for 100% viability.

### B3Z T Cell Activation

1.5×10^4^ MC38-OVA cells were seeded in a treated 96 well plate, either with or without compounds (BIIB021 and **CliMB-325)** at indicated concentrations at 37°C. After 48 hours, the culture media was replaced with media containing 10^5^ B3Z cells, which were co-incubated with the MC38-OVA cells for 6 hours. Cells were centrifuged (1100 x g, 5 min) in a Thermo Scientific Jouan C4i centrifuge and the supernatant was removed. Lysis buffer containing 0.2% saponin, 500 mM CPRG reagent, 20 mM MgCl_2_, and 100 mM β-mercaptoethanol in 1X PBS was added to each well. After 45 minutes, absorbance at 570 nm was recorded using a BioTek Synergy H1 Microplate Reader.

### Molecular Docking Studies

Conformational predictions of **CliMB-325** in Hsp90 were performed using RosettaLigand using the crystal structure of BIIB021 bound to Hsp90 (PDB ID: 3qdd).^63–66^ Native crystal structure was prepared for docking by removing all water molecules and co-crystallized ligands. PyMOL was used for visualization of the docking results.

### Mild Acid Elution (MAE) of MHC-I-Bound Peptides

The MAE protocol was adapted from a previously published protocol.^67^ MAEs from CT26 cells were performed with 1 x 10^7^ CT26 cells per sample. Cells were washed three times with PBS then were treated for 90 seconds with MAE buffer. The MAE buffer consisted of 131 mM citric acid, 66 mM Na_2_HPO_4_, and 150 mM NaCl adjusted to pH 3.3 with NaOH. Following treatment with the MAE buffer, the eluted peptide solution was centrifuged (4000 x g, 5 min) in a Thermo Scientific Jouan C4i centrifuge. The peptide-containing supernatant was collected and acidified with 0.1% TFA. The obtained peptide solution was further purified on Oasis HLB columns (barrel size 1 cm^3^, 30 mg of sorbent; Waters, product no.: WAT094225) prerinsed with 100% acetonitrile (MeCN)/0.1% TFA. After equilibration with 100% H_2_O/0.1% TFA, sample loading, and washing with 95% MeCN/0.1% TFA, peptides were eluted with 60% MeCN/0.1% TFA. The eluate was filtered through a 15 mL Amicon ultrafilter device with 3 kDa molecular weight cutoff (Merck Millipore, Cat.-No. UFC901024) then lyophilized to dryness using a Labconco Freezone 4.5 L (−84°C) lyophilizer.

### Liquid Chromatography

Peptide separation was carried out on a Vanquish Neo UHPLC system using a trap-and-elute setup. An Aurora Frontier™ TS C18 column (IonOpticks; 60 cm × 75 µm, 1.7 µm particles) was used for peptide separation. The mobile phases used were: Phase A — 0.1% formic acid (FA) in water; Phase B — 80% acetonitrile (ACN) with 0.1% FA in water. A 40-minute gradient was applied at 0.3 µl/min: 15% to 50% MPB from 0.1 to 40 min, followed by 50% to 99% mobile phase buffer B (MPB) from 40.1 to 42 min, held at 99% MPB until 50 min, and re-equilibrated at 1% MPB for 35 minutes.

### Mass Spectrometry Data Acquisition

Data were acquired using an Orbitrap Astral mass spectrometer in a DDA mode. For MS1 scans, the Orbitrap resolution was set at 120,000 with an AGC target of 100%. MS1 spectra were recorded over an m/z range of 350–1350, with a maximum injection time of 50 ms. The isolation window for MS2 precursor selection was set to 1.2 m/z. Up to 30 scans were acquired per MS1 cycle for precursor ions with intensities greater than 5.0 x 10^3^ and charge states ranging from 2 to 6. MS2 fragmentation was performed with HCD at 25% collision energy and a maximum injection time of 25 ms. Dynamic exclusion was turned on with a duration of 20 seconds.

### DDA Data Analysis

Spectra were converted to mzXML using a modified version of ReAdW.exe. Mass spectra were processed using a COMET-based software pipeline and searched against the mouse UniProt database (downloaded on October 3^rd^, 2024). Database searches were performed using a precursor ion tolerance of 50-ppm and 0.02 Da fragment ion tolerance. No enzyme specificity was specified for peptide identification. Carbamidomethylation of cysteine residues (+57.021 Da) were set as static modifications, while oxidation of methionine residues (+15.995 Da) was set as a variable modification. Peptide-spectrum matches (PSMs) were adjusted to a 1% false discovery rate (FDR) using standard target-decoy approaches.^68^ Sequenced peptides were further filtered based on length (8–14 amino acids) to ensure likelihood of MHC-I binding. The obtained peptide list was input into NetMHCpan 4.1 database to predict binding compatibility with the MHC-I allotypes expressed by CT26 cells (H-2-Dd, H-2-Kd, and H-2-Ld).^69^ Prediction score was reported by NetMHCpan 4.1, and downstream peptide analysis was only performed on peptides which were modeled to bind with a prediction score of 2.0 or lower and thus expected to be putative binders of MHC-I molecules.

### Synthesis of Target Compounds

NMR spectroscopy was performed in DMSO-*d*_6_ on a Varian 600 MHz spectrophotometer. Chemical shifts are reported in ppm (δ) and coupling constants (J) are reported in Hertz [Hz]. High-resolution electrospray ionization mass spectrometry (HR-ESI-MS) was performed using an Agilent 1260 Infinity II Prime liquid chromatography (LC) system. Compounds were analyzed for purity using RP analytical HPLC equipped with Waters 1525 with a 2489 UV/Visible Detector monitoring at 311 nm wavelength, on a Phenomenex Luna 5 µM C18(2) 250 x 4.6 mm column using gradient elution. Flow rate = 1.0 mL/min; mobile phase A = 0.1% trifluoroacetic acid (TFA) (v/v) in water; mobile phase B = 0.1% TFA in methanol (MeOH); gradient: 5% B for 5 min, 5– 100% B for 20 min, 100% B hold for 3 min, then 100–5% B for 15 min. All compounds are >95% pure by HPLC analysis

## 9-propargyl-2-amino-6-chloropurine

9-propargyl-2-amino-6-chloropurine was synthesized based on literature procedure.^70^ 2-amino-6-chloropurine (3.0 g, 1 eq) was suspended DMF (50 mL) followed by addition of anhydrous K_2_CO_3_ (2.934 g, 1.2 eq) and stirring under N_2_ atmosphere for 1 hour. After this time, propargyl bromide (1.894 g, 0.9 eq) was added and stirred for 48 hours under N_2_ atmosphere at room temperature. DMF was evaporated at 60°C under high vacuum to afford a yellowish-white powder. A 1:2 ratio of minor and major compound was produced as determined by NMR. The crude material was purified by reverse-phase preparative high-performance liquid chromatography (RP-HPLC) equipped with Waters 1525 with a 2489 UV/Visible Detector monitoring at 311 nm wavelength, on a Phenomenex Luna Omega 5 µM Polar C18 250 x 21.2 mm column using gradient elution with using H_2_O/MeOH with 0.1% TFA at 10 mL/min. The HPLC fractions of the major compound were concentrated under reduced pressure using a rotary evaporator, then lyophilized to dryness using a Labconco Freezone 4.5 L (−84°C) lyophilizer and characterized by NMR which matched with the reported compound.^71^ This product was analyzed for purity using RP analytical HPLC equipped with Waters 1525 with a 2489 UV/Visible Detector monitoring at 311 nm wavelength, on a Phenomenex Luna 5 µM C18(2) 250 x mm column using gradient elution with using H_2_O/MeOH with 0.1% TFA at 1 mL/min. The major product was used for click chemistry. White solid;^71^ ^1^H NMR (600 MHz, DMSO-*d_6_)* δ 8.18 (s, 1H, 8-H), 7.02 (brs, 2H, -NH_2_), 4.93 (d, 2H, J=1 Hz, -CH_2_), 3.48 (t, 1H, J = 1 Hz, C≡CH). HRMS: m/z calculated for C_8_H_6_ClN_5_ [M+H]^+^ 208.0385, found 208.0388.

### 380 Compound Library of Triazole-Containing BIIB021 Derivatives

Triazole analogs were synthesized based on literature procedure.^72^ Azide solutions from the azide library in Plates 1-5 were initially at a concentration of 100 mM in DMSO. Azides were added in each well of a 96-well plate at a concentration of 10 mM. To each well of this newly loaded plate, L-ascorbic acid solution was added to a concentration of 40 mM along with 10 mM of 9-propargyl-2-amino-6-chloropurine (synthesis shown in *Scheme S1*) and 2 mM of CuSO_4_/THPTA in a solution of DMSO and water at a 3:2 ratio to a total volume of 100 µL. The plates were sealed and swirled at 250 rpm and 37°C for 20 hours to afford the corresponding triazole product in each well.

### CliMB-325 (2-(4-((2-amino-6-chloro-9H-purin-9-yl)methyl)-1H-1,2,3-triazol-1-yl)-5-chlorobenzonitrile.)

In a 50 mL conical tube, the following reagents were added: 10 mM of 2-azido-5-chlorobenzonitrile (azide #325 from 380 compound screen), 40 mM of aqueous L-ascorbic acid, 10 mM of 9-propargyl-2-amino-6-chloropurine (synthesis shown in *Scheme S1*), 2 mM of aqueous CuSO_4_/THPTA solution, in a 3:2 ratio of DMSO to water at a total volume of 15 mL. The tube was swirled at 250 rpm and 37°C for 20 hours to yield **CliMB-325**. The compound was purified by reverse-phase preparative high-performance liquid chromatography (RP-HPLC) equipped with Waters 1525 with a 2489 UV/Visible Detector monitoring at 311 nm wavelength, on a Phenomenex Luna Omega 5 µM Polar C18 250 x 21.2 mm column using gradient elution with using H_2_O/MeCN with 0.1% TFA at 10 mL/min. The HPLC fractions of the desired purified product were concentrated under reduced pressure using a rotary evaporator, then lyophilized to dryness using a Labconco Freezone 4.5 L (−84°C) lyophilizer. The product was analyzed for purity using RP analytical HPLC equipped with Waters 1525 with a 2489 UV/Visible Detector monitoring at 311 nm wavelength, on a Phenomenex Luna 5 µM C18(2) 250 x mm column using gradient elution with using H_2_O/MeCN with 0.1% TFA at 1 mL/min. The final product was stored at −20°C until further use, and stocks were made at 10 mM in DMSO. White solid; ^1^H NMR (600 MHz, DMSO-*d_6_)* d 8.73 (s, 1H, 8-H), 8.36 (d, 1H, J= 1 Hz, Ar-H), 8.24 (s, 1H, triazole-H), 8.04 (dd, 1H, J= 3.6 Hz, Ar-H), 7.88 (d, 1H, J= 3.6 Hz, Ar-H), 6.95 (brs, 2H, -NH), 5.51 (s, 2H, -CH_2_); ^13^C NMR (150 MHz, DMSO-*d_6_*) d 159.9, 153.9, 149.4, 143.4, 142.9, 136.5, 134.7, 134.5, 134.1, 127.3, 124.7, 123.2, 114. 6, 108.6, 38.0. HRMS: m/z calculated for C_15_H_10_Cl_2_N_9_ [M+H]^+^ 386.0431, found 386.0439.

## SUPPORTING INFORMATION

- Additional figures, tables, and materials/methods are included in the Supporting Information File (PDF)
- Molecular formula strings (CSV)
- PDB file for BIIB021 bound to Hsp90 (PDB)
- PDB file for **CliMB-325** bound to Hsp90 (PDB)
- Table of MHC-I peptide sequences obtained by MS (XLSX)

## AUTHOR INFORMATION

### Notes

The authors declare no competing financial interest.

## ACKNOWLEDGEMENTS

This study was supported by the NIH grant R35GM124893 (M.M.P.)

## ABBREVIATIONS USED

APC: allophycocyanin
BET: bromodomain and extra-terminal
CPRG: chlorophenol red-β-galactopyranoside
CuAAC: Cu(I)-catalyzed azide-alkyne cycloaddition
DNMT: DNA methyltransferase
IMiD: immunomodulatory imide drug
IL-2: interleukin-2
MFI: mean fluorescence intensity
MAE: mild acid elution
OVA: ovalbumin
pMHC: peptide-MHC complex
PD-1: programmed cell death protein 1
PD-L1: programed death-ligand 1
STING: stimulator of interferon genes
TCR: T cell receptor

## TABLE OF CONTENTS GRAPHIC

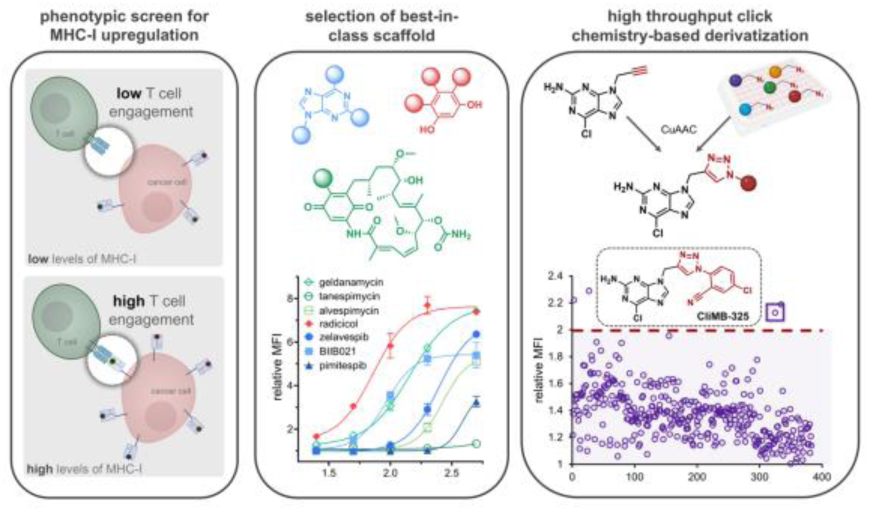

## Notes

### Competing Interest Statement

The authors have declared no competing interest.

### Summary of Updates

Title updated to emphasize the click chemistry-based discovery approach; author affiliations updated; abstract updated for conciseness; Figure 1 revised to include flow chart; Figure 4C updated to include docking image of BIIB021 into Hsp90; Figure 4E updated to include revised MTT viability assay data; new section appended in results to include LC-MS/MS identification of MHC-I peptides; Supplemental files updated

