## Supplementary Information for "Small Molecule Modulation of MHC-I Surface Expression: A Click Chemistry-Based Discovery Approach"

### Table of Contents


| | |
|---|---|
| **SUPPLEMENTARY FIGURES** | **S3-S15** |
| Figure S1. *Chemical structures for 25 compound library* | S3 |
| Figure S2. *Flow cytometry analysis of CT26 cells treated with 25-member library at 5 μM* | S4 |
| Figure S3. *Analytical HPLC of reaction between 9-propargyl-2-amino-6-chloropurine and 2-azido-1-(4-methoxy-phenyl)-ethanone* | S5 |
| Figure S4. *Analytical HPLC of reaction between 9-propargyl-2-amino-6-chloropurine and 4-azido-1-butanamine* | S6 |
| Figure S5. *Analytical HPLC of reaction between 9-propargyl-2-amino-6-chloropurine and Boc-4-azido-L-phenylalanine* | S7 |
| Figure S6. *Chemical structures for 380 compound library* | S8 |
| Figure S7. *Chemical structures of triazole products from azides 1-8* | S9 |
| Figure S8. *Flow cytometry analysis of CT26 cells treated with 1 μM 9-propargyl-2-amino-6-chloropurine or a 1:10,000 dilution of CuAAC click reagents* | S12 |
| Figure S9. *Flow cytometry analysis of CT26 cells treated with triazole products of 3, 27, 325, and 335* | S13 |
| Figure S10. *Flow cytometry analysis of CT26 cells treated with **CliMB-325** regioisomer* | S14 |
| Figure S11. *Length distribution of MHC-I peptides isolated by MAE from CT26 cells* | S15 |
| **MATERIALS AND METHODS** | **S16-S18** |
| **Materials** | S16 |
| **Experimental Methods** | S16-18 |
| Mammalian Cell Culture | S16 |
| Flow Cytometry-Based Assays | S16 |
| MTT Cell Viability Assay | S16 |
| B3Z T Cell Activation | S17 |
| Molecular Docking Studies | S17 |
| Mild Acid Elution (MAE) of MHC-I-Bound Peptides | S17 |
| Liquid Chromatography | S17 |
| Mass Spectrometry Data Acquisition | S18 |
| DDA Data Analysis | S18 |
| **SYNTHESIS AND CHARACTERIZATION** | **S19-27** |
| Scheme S1: *Synthesis of 9-propargyl-2-amino-6-chloropurine* | S19 |
| Scheme S2: *High-Throughput Synthesis of Triazole-Containing BIIB021 Derivatives* | S22 |
| Scheme S3: *Synthesis of **CliMB-325*** | S23 |
| **REFERENCES** | **S28-29** |

#### SUPPLEMENTARY FIGURES

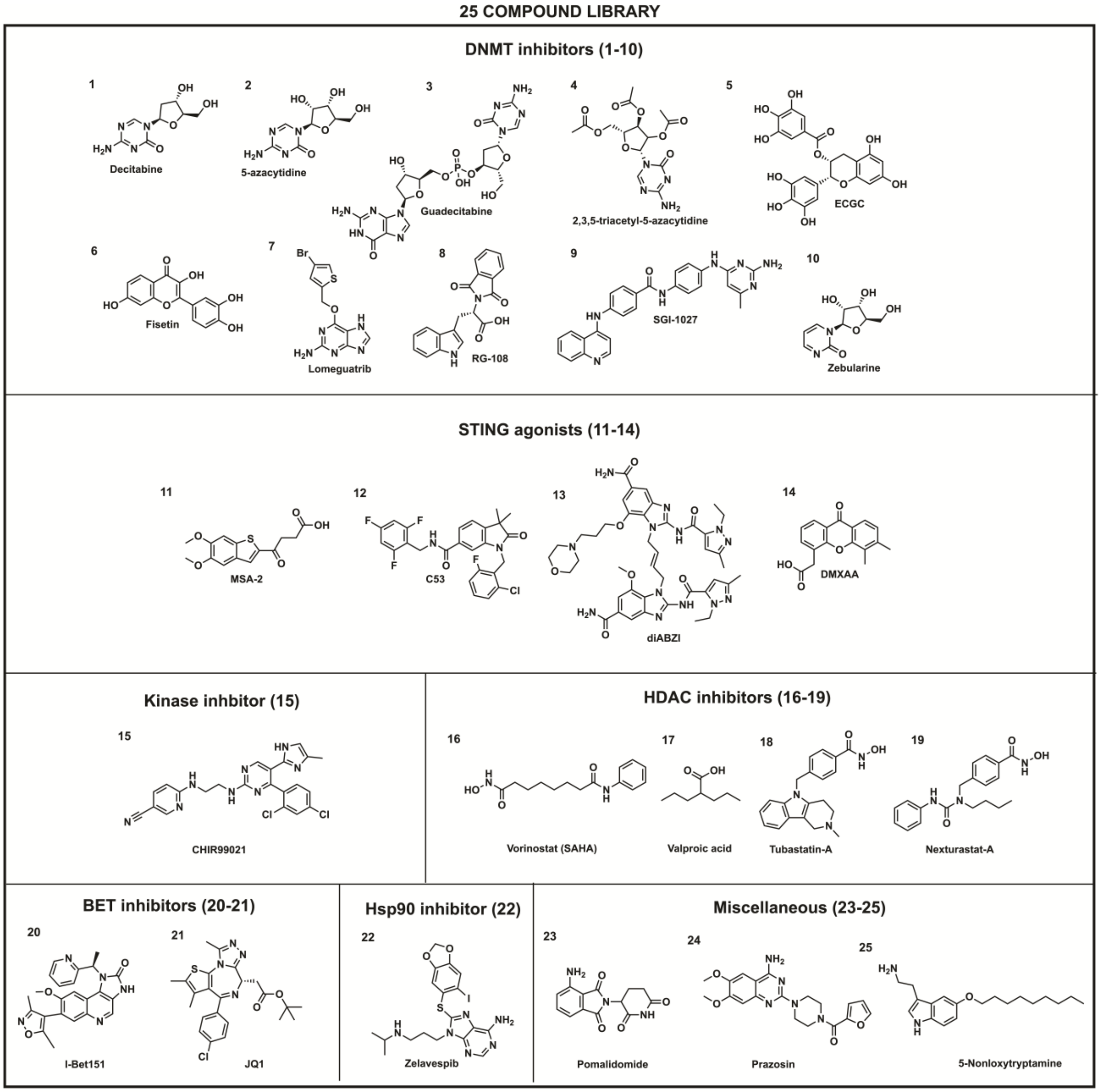


**Figure S1.** Structures of 25 compounds in library of small molecule MHC-I inducers, subdivided by class. Compounds 1-10 are DNMT inhibitors, 11-14 are STING agonists, 15 is a kinase inhibitor, 16-19 are HDAC inhibitors, 20-21 are BET inhibitors, 22 is an Hsp90 inhibitor, and 23-25 are miscellaneous compounds.

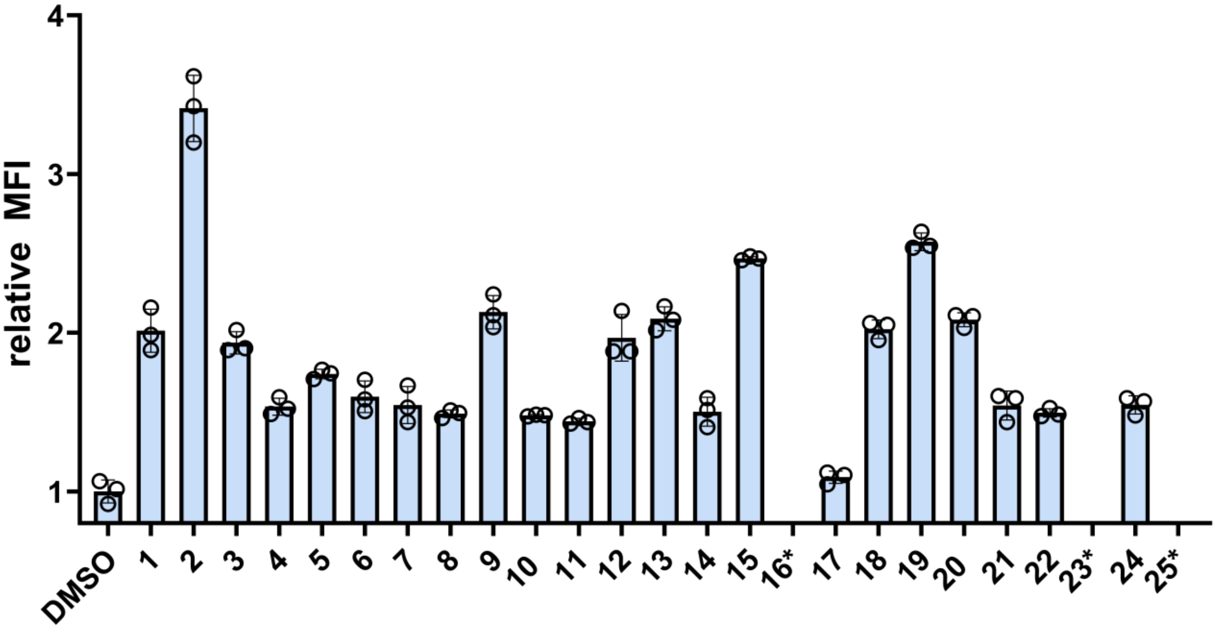


**Figure S2.** Flow cytometry analysis of CT26 cells treated with 25-member library at 5 μM. H-2K$^d$ surface expression was measured by APC-conjugated anti-mouse H-2K$^d$ antibody. MFI means fluorescence intensity of the level of fluorescence relative to the DMSO control. Compounds containing no data bar and denoted with * indicate that the compound was toxic to the cells at 5 μM concentration. Data are represented as mean ± SD (*n*=3).

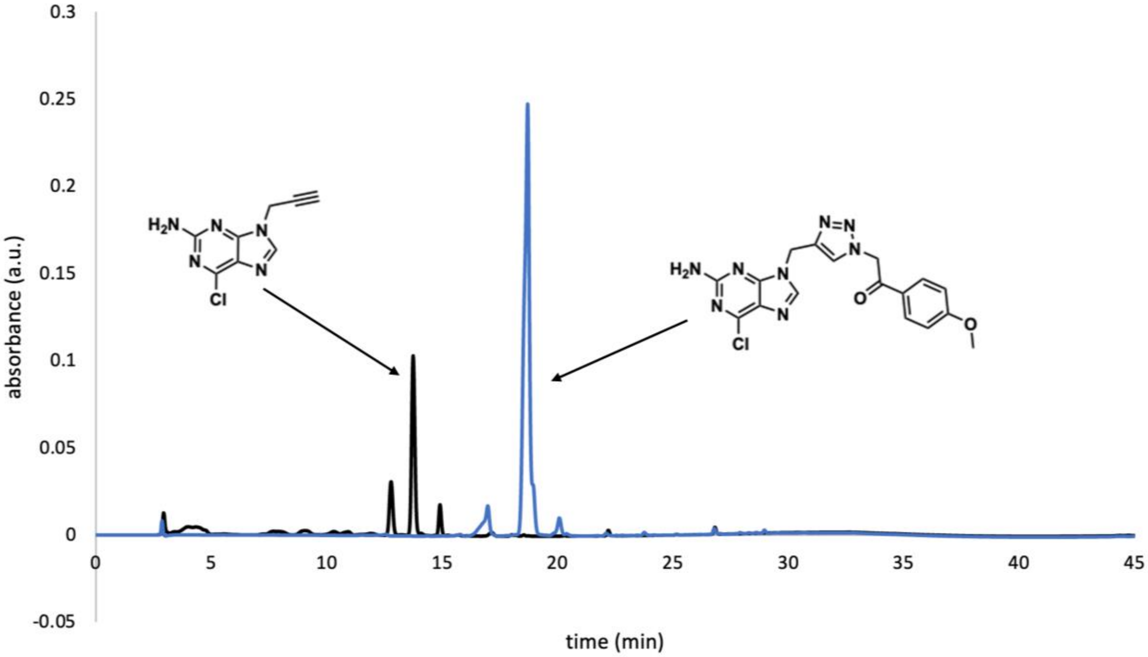


**Figure S3**. Analytical HPLC of reaction between 9-propargyl-2-amino-6-chloropurine and 2-azido-1-(4-methoxy-phenyl)-ethanone. 9-propargyl-2-amino-6-chloropurine and 2-azido-1-(4-methoxy-phenyl)-ethanone, each at a concentration of 10 mM, were reacted with 40 mM L-ascorbic acid and 2 mM $CuSO_4$/THPTA in a 3:2 ratio of DMSO to water, with a total reaction volume of 100 µL. Overlaid HPLC chromatograms of reaction mixture prior to addition of 2-azido-1-(4-methoxy-phenyl)-ethanone (black) and the full reaction mixture after incubation shaking at 37 $^{o}$C for 20 h (blue) are shown.

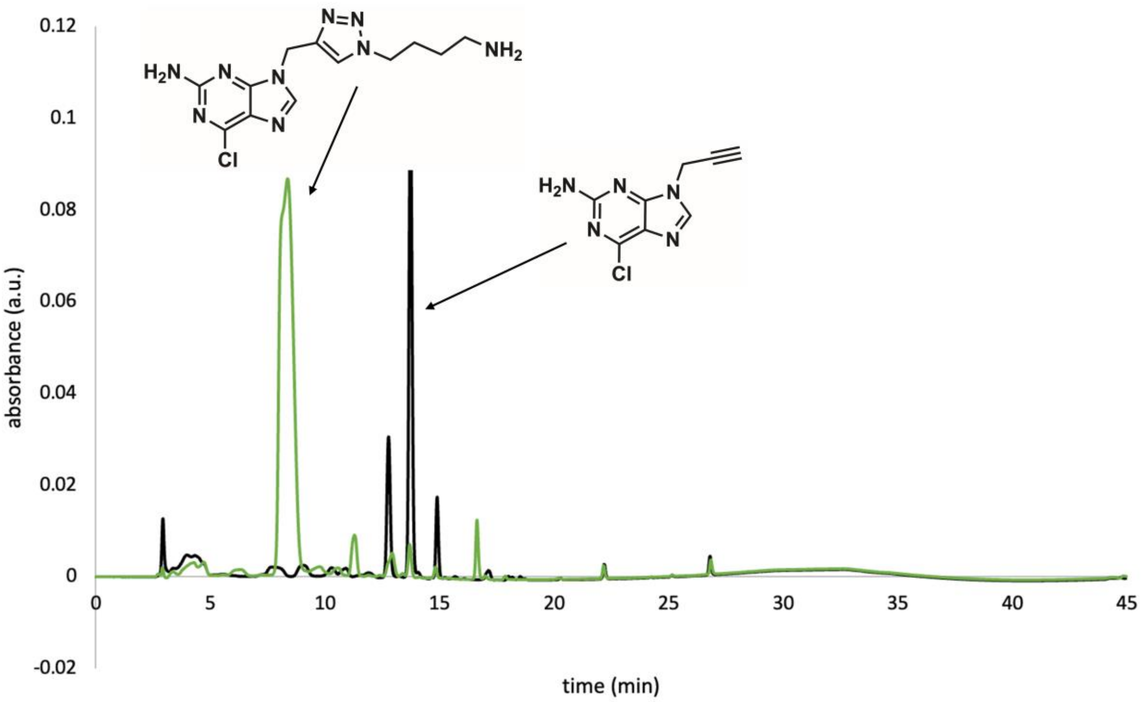


**Figure S4.** Analytical HPLC of reaction between 9-propargyl-2-amino-6-chloropurine and 4-azido-1-butanamine. 9-propargyl-2-amino-6-chloropurine and 4-azido-1-butanamine, each at a concentration of 10 mM, were reacted with 40 mM L-ascorbic acid and 2 mM $CuSO_4$/THPTA in a 3:2 ratio of DMSO to water, with a total reaction volume of 100 µL. Overlaid HPLC chromatograms of reaction mixture prior to addition of 4-azido-1-butanamine (black) and the full reaction mixture after incubation shaking at 37 $^{o}$C for 20 h (green) are shown.

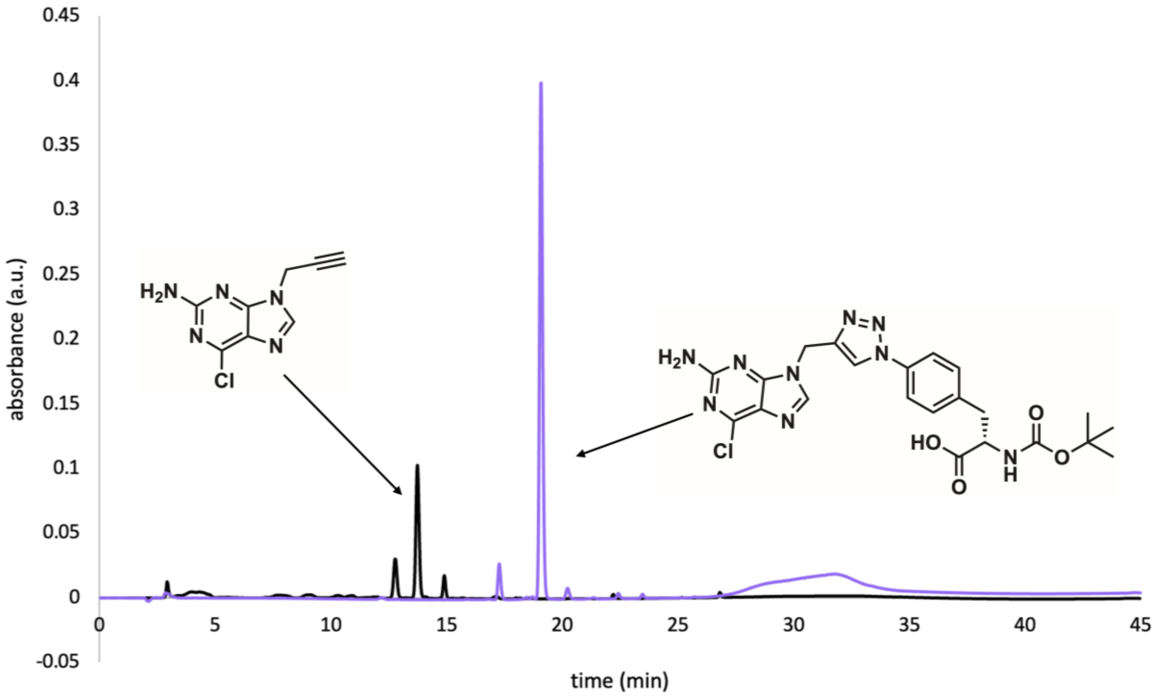


**Figure S5.** Analytical HPLC of reaction between 9-propargyl-2-amino-6-chloropurine and Boc-4-azido-L-phenylalanine. 9-propargyl-2-amino-6-chloropurine and Boc-4-azido-L-phenylalanine, each at a concentration of 10 mM, were reacted with 40 mM L-ascorbic acid and 2 mM $CuSO_4$/THPTA in a 3:2 ratio of DMSO to water, with a total reaction volume of 100 µL. Overlaid HPLC chromatograms of reaction mixture prior to addition of Boc-4-azido-L-phenylalanine (black) and the full reaction mixture after incubation shaking at 37 $^{o}$C for 20 h (purple) are shown.

1
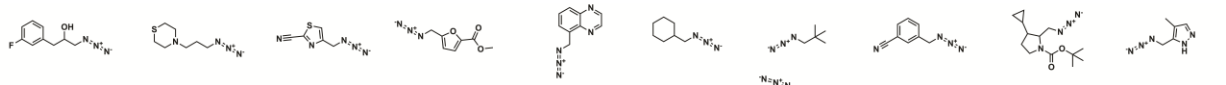

11
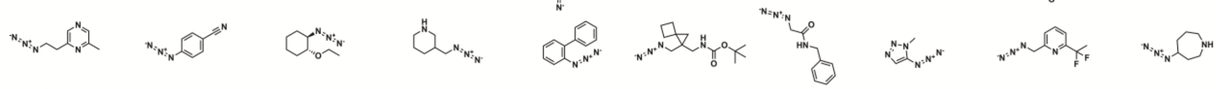

21
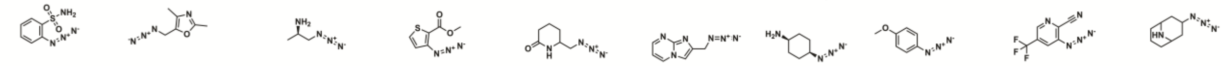

31
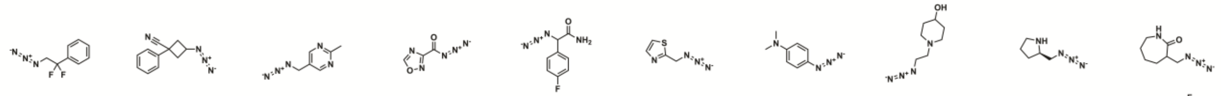

41
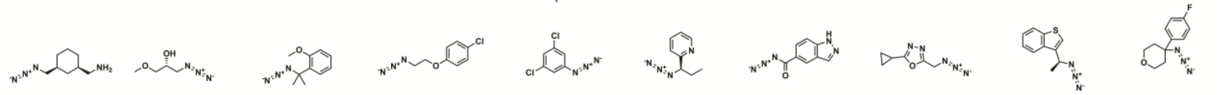

51
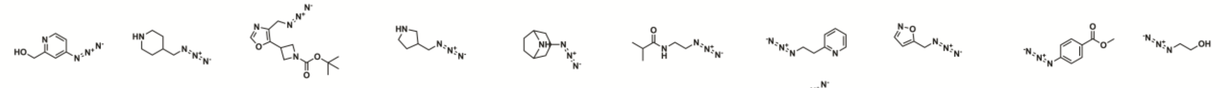

61
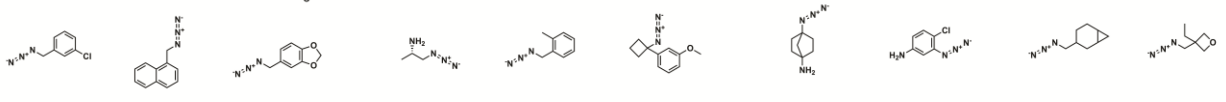

71
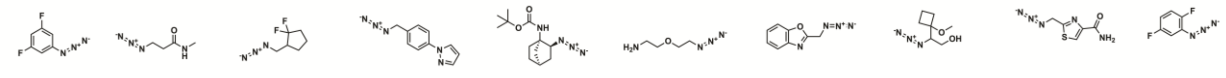

81
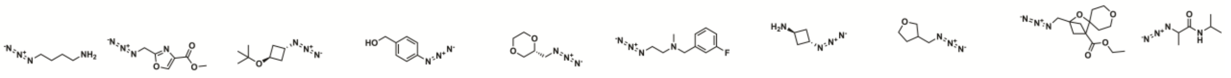

91
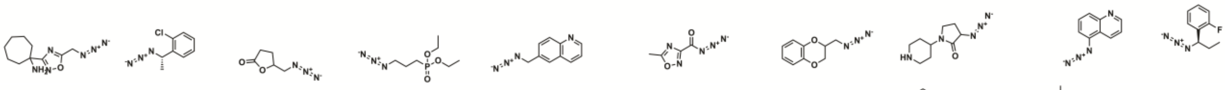

101
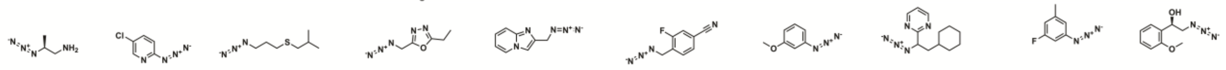

111
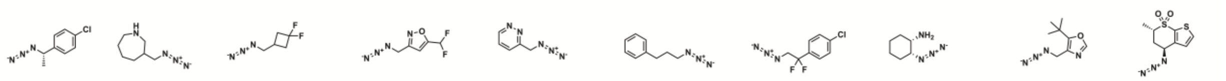

121
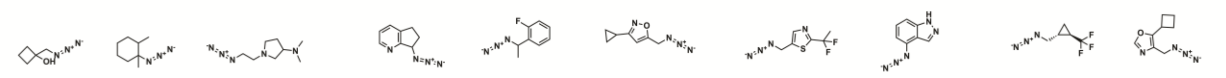

131
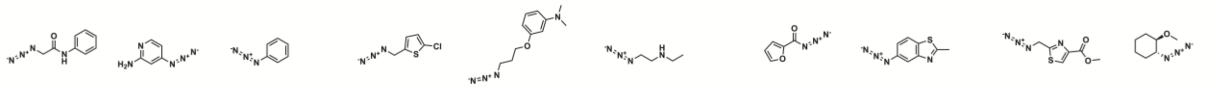

141
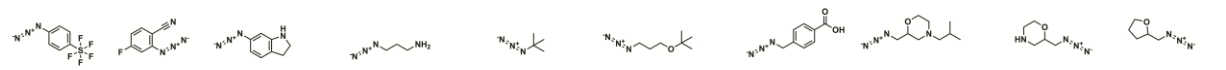

151
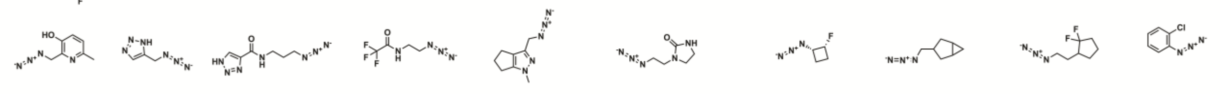

161
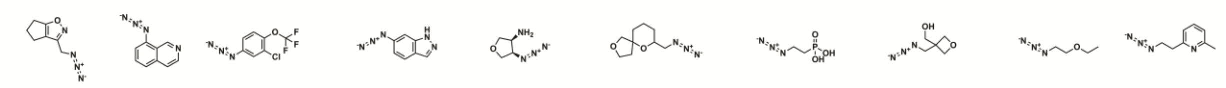

171
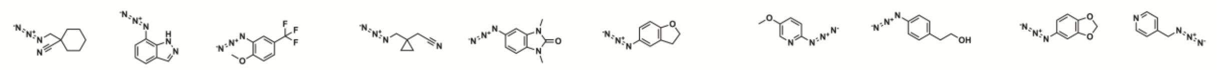

181
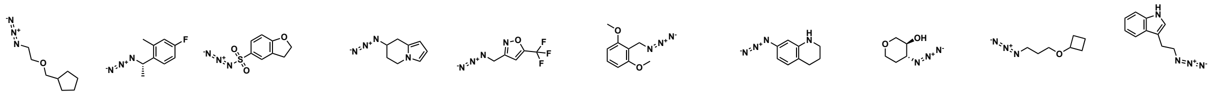

191
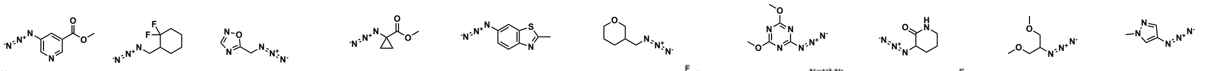

201
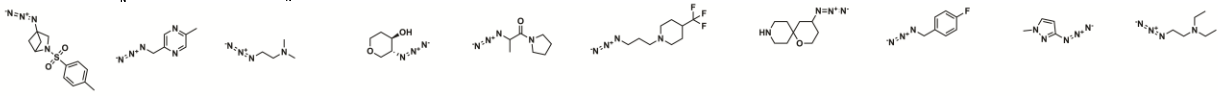

211
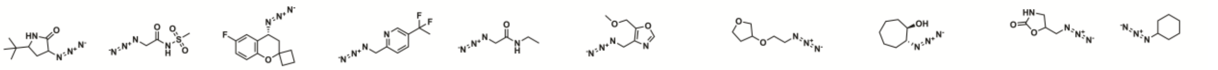

221
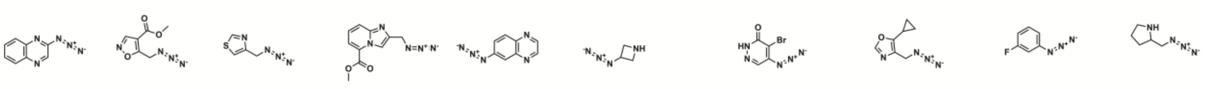

231
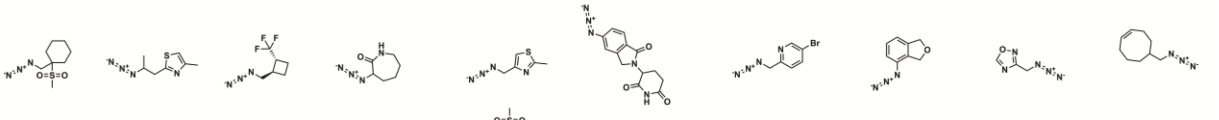

241
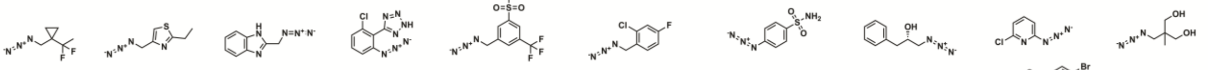

251
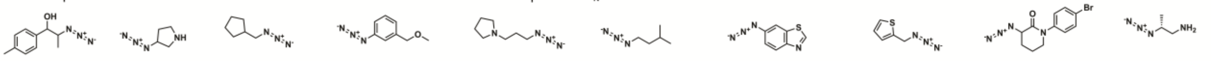

261
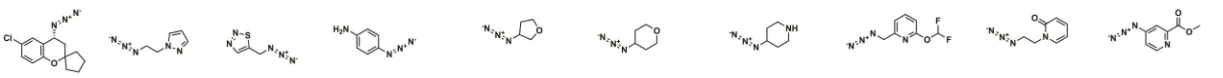

271
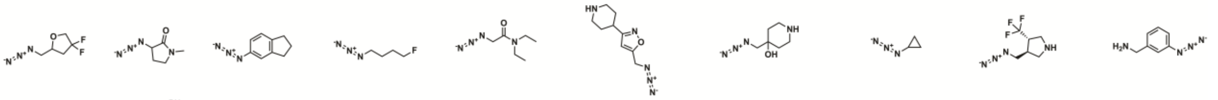

281
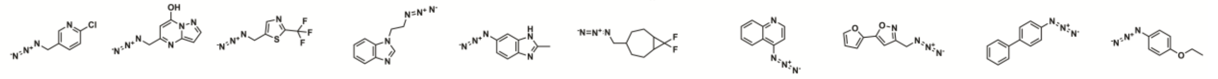

291
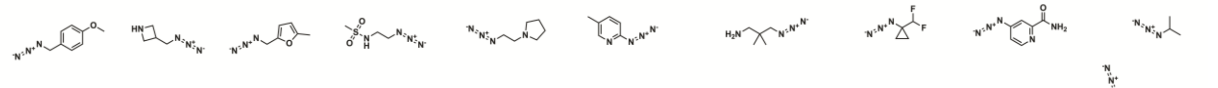

301
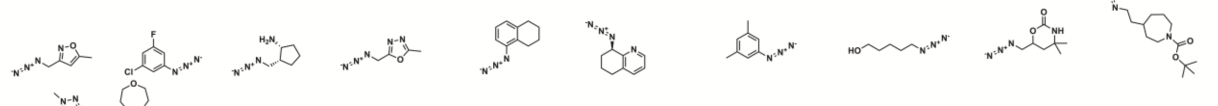

311
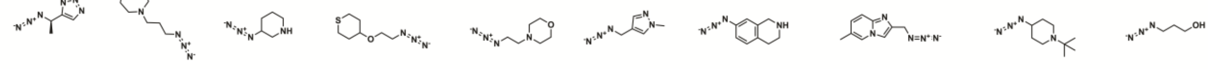

321
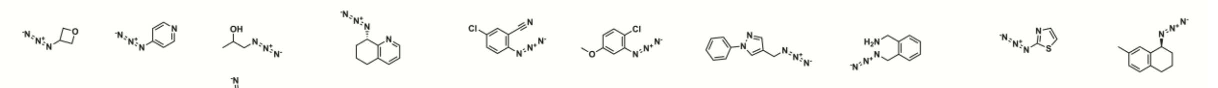

331
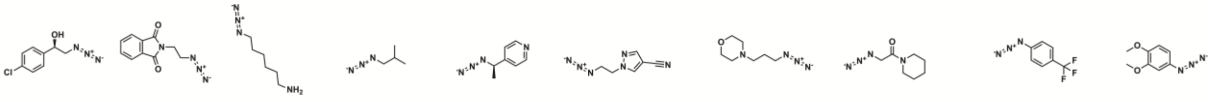

341
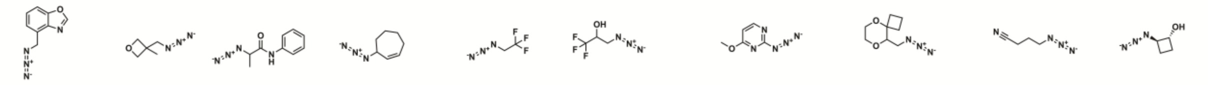

351
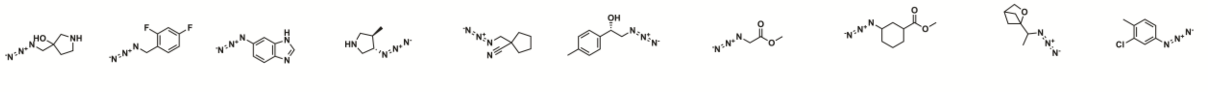

361

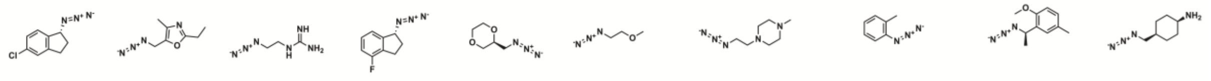

371

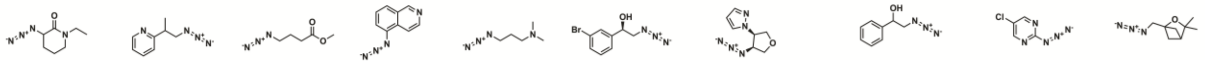

**Figure S6.** Structures of azide-containing small molecules from 380 compound library.

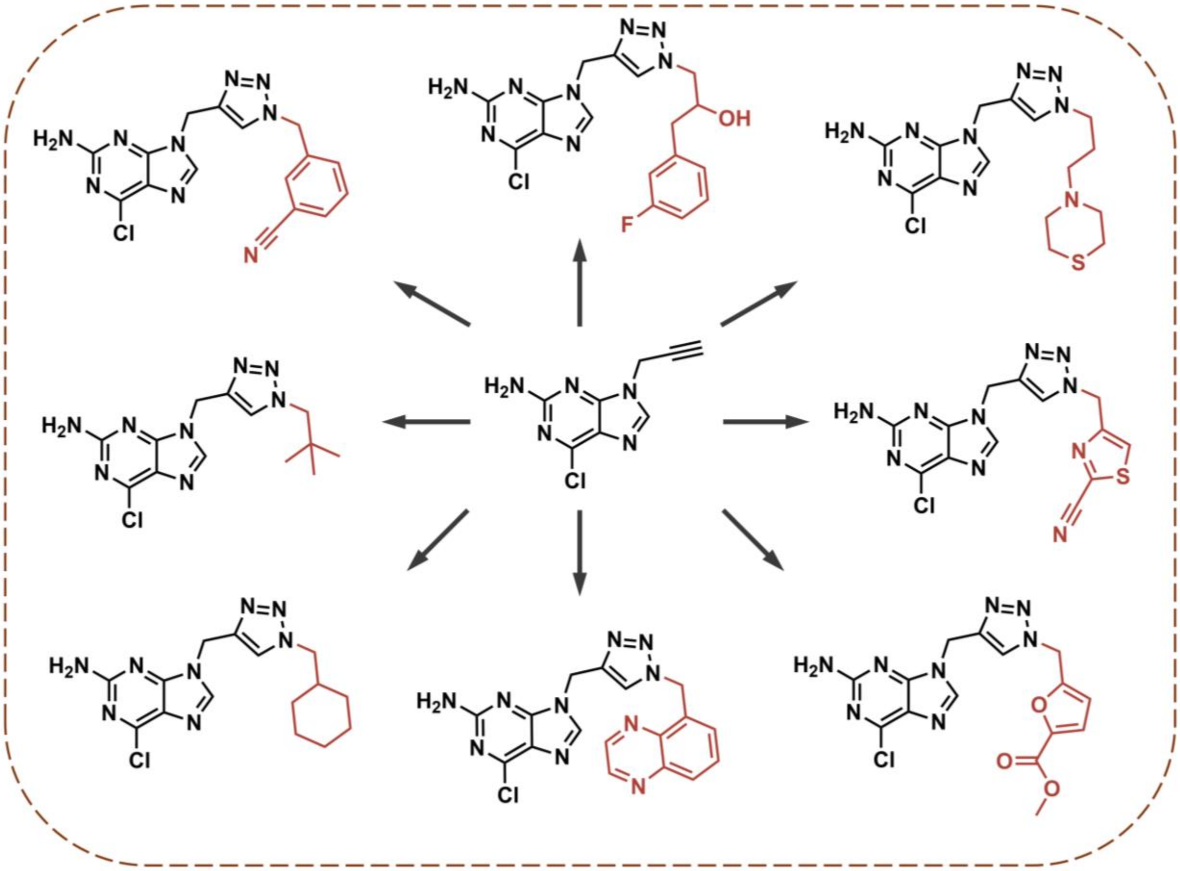

**Figure S7.** Chemical structures of triazole products formed from click reaction between 9-propargyl-2-amino-6-chloropurine and azide compounds 1-8 of 380 library.

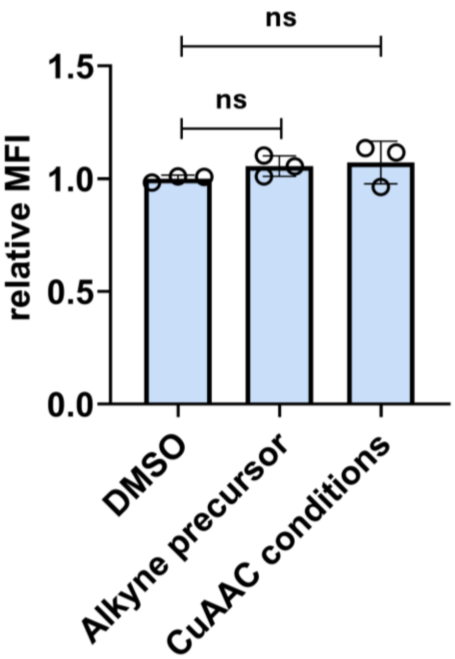


**Figure S8.** Flow cytometry analysis of CT26 cells treated with 1 μM 9-propargyl-2-amino-6-chloropurine or a 1:10,000 dilution of CuAAC click reagents (dilution used for Figure 4B). H-2K$^d$ expression was measured by APC anti-mouse H-2K$^d$ antibody. MFI is mean fluorescence intensity of the level of fluorescence relative to the DMSO control. Data are represented as mean ± SD (n=3). P-values were determined by a two-tailed *t*-test (ns = not significant).

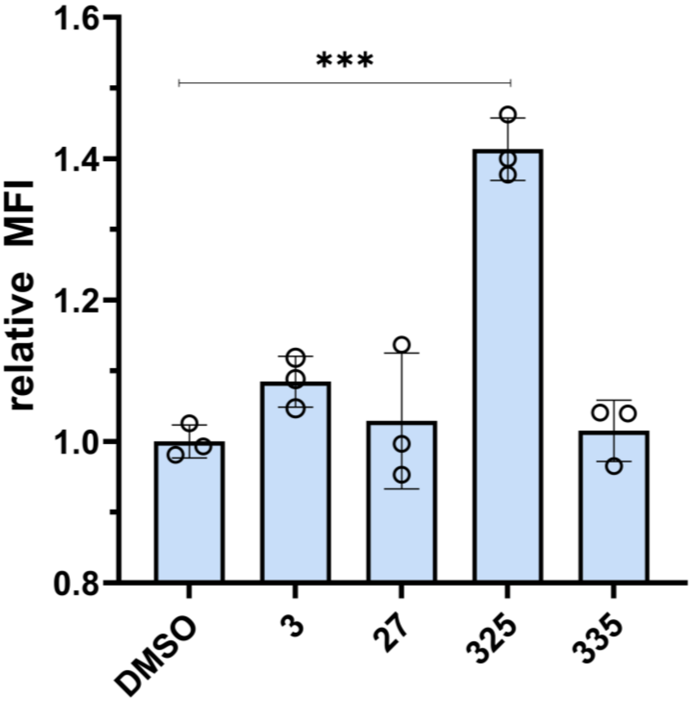


**Figure S9.** Flow cytometry analysis of CT26 cells treated with 1:20,000 dilution of click reaction (500 nM assuming complete conversion) mixtures containing azides 3, 27, 325, and 335. H-2K$^d$ expression was measured by APC anti-mouse H-2K$^d$ antibody. MFI means fluorescence intensity of the level of fluorescence relative to the DMSO control. Data are represented as mean ± SD (n=3). p-values were determined by a two-tailed *t*-test (*** $p < 0.001$).

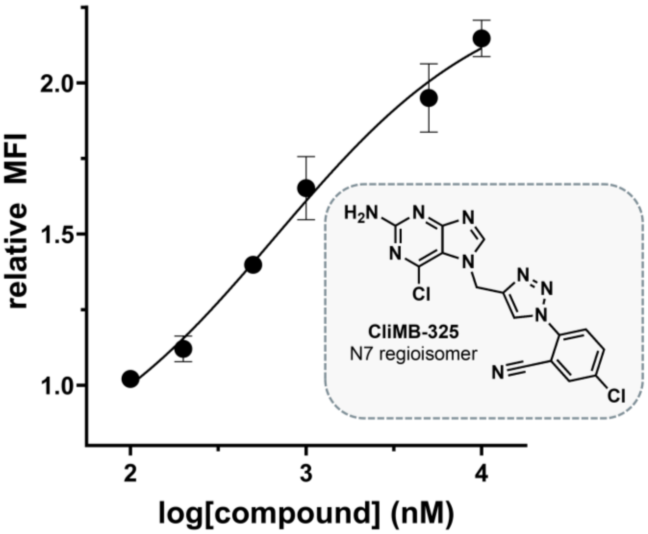


**Figure S10.** Dose-response curve and chemical structure of the regioisomer of **CliMB-325,** formed from a reaction between the minor N7 regioisomer of the alkyne-modified BIIB021 precursor (7-propargyl-2-amino-6-chloropurine) and 2-azido-5-chlorobenzonitrile (azide 325 from 380 compound screen). CT26 cells were treated with varying concentrations of the **CliMB-325** regioisomer. H-2$K^d$ expression was measured by APC anti-mouse H-2$K^d$ antibody via flow cytometry. MFI means fluorescence intensity of the level of fluorescence relative to the DMSO control. Data are represented as mean ± SD (n=3), and Boltzmann sigmoidal curves were fitted to the data using GraphPad Prism. $EC_{50}$ values are the concentration of compound needed to achieve 50% of the maximal MHC-I surface expression levels.

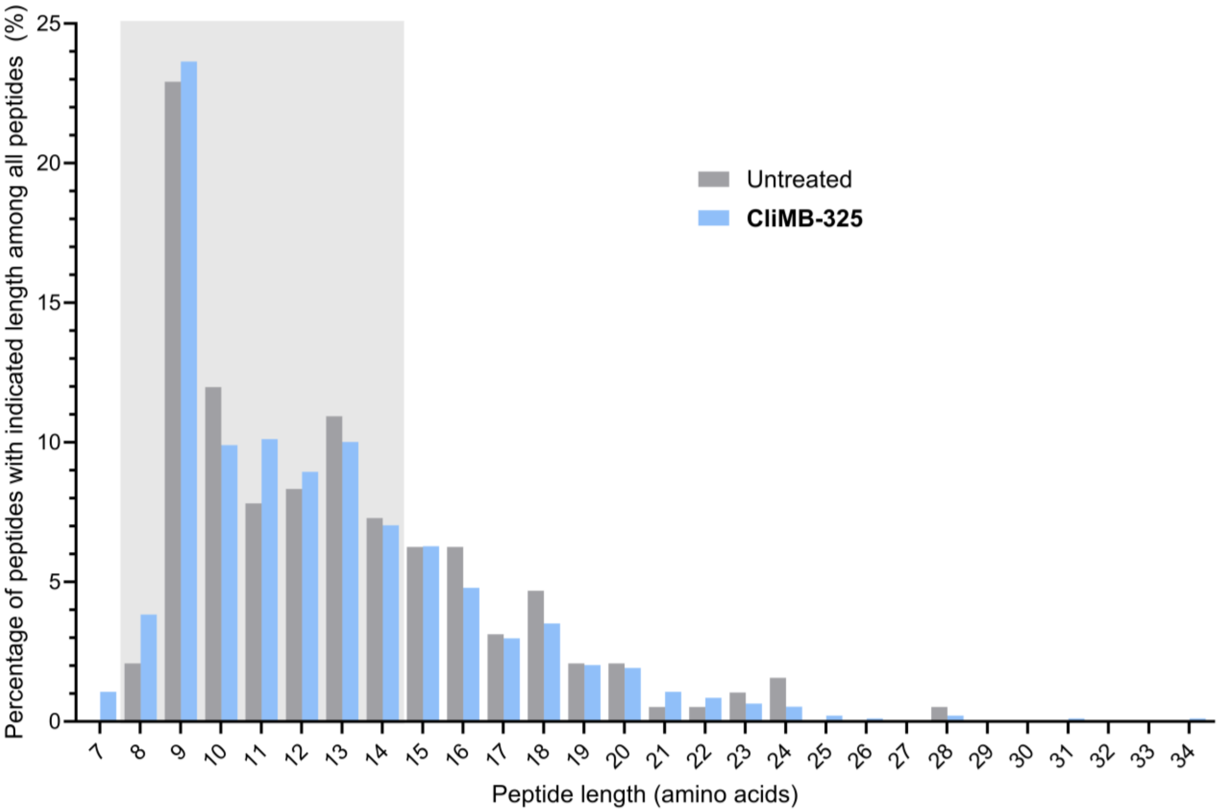


**Figure S11.** Length distribution of MHC-I peptides isolated by mild acid elution (MAE) from CT26 cells treated with (blue) or without (gray) **CliMB-325**. Data in displayed histograms are derived from a single cell culture, and peptides were obtained from 1 x $10^7$ cells per sample. Light gray shaded area indicates the *in silico* length filter of 8 to 14 amino acids that was applied for all analyses (except for this figure). The summed fraction of 8- to 14-mers among all peptides was 64% and 66% for untreated and **CliMB-325**-treated samples, respectively.

#### MATERIALS

**Reagents.** All library compounds were purchased from either Selleck Chemicals, AK Scientific, A2B Chem, MedChemExpress, GlpBio, Cayman Chemical Company, or AmBeed. Compounds were solubilized in DMSO and stored at -20°C. Recombinant murine and human IFN-γ were purchased from PeproTech. APC-labeled antibodies against H-2K$^d$/H-2D$^d$, HLA-A,B,C, and H-2K$^b$ bound to SIINFEKL were purchased from BioLegend. The library of 380 azide-containing small molecules were purchased from Enamine (catalog # AZD-380-X-100). For the synthesis of **CliMB-325**, 2-amino-6-chloropurine was purchased from AmBeed (catalog # A135577) and 2-azido-5-chlorobenzonitrile was purchased from Enamine (catalog # EN300-279694). Dulbecco's Modified Eagle's Medium (DMEM), Roswell Park Memorial Institute (RPMI) 1640 medium, and McCoy's 5A medium were purchased from VWR. Fetal Bovine Serum (FBS) and penicillin-streptomycin were purchased from Sigma-Aldrich.

#### SYNTHESIS AND CHARACTERIZATION

##### Scheme S1. Synthesis of 9-propargyl-2-amino-6-chloropurine

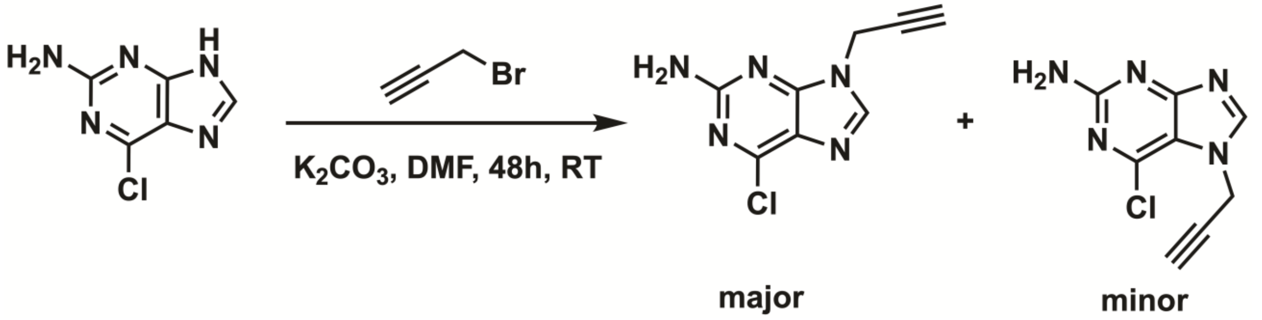


9-propargyl-2-amino-6-chloropurine was synthesized based on literature procedure.[8] 2-amino-6-chloropurine (3.0 g, 1 eq) was suspended DMF (50 mL) followed by addition of anhydrous $K_2CO_3$ (2.934 g, 1.2 eq) and stirring under $N_2$ atmosphere for 1 hour. After this time, propargyl bromide (1.894 g, 0.9 eq) was added and stirred for 48 hours under $N_2$ atmosphere at room temperature. DMF was evaporated at 60°C under high vacuum to afford a yellowish-white powder. A 1:2 ratio of minor and major compound was produced as determined by NMR. The crude material was purified by reverse-phase preparative high-performance liquid chromatography (RP-HPLC) equipped with Waters 1525 with a 2489 UV/Visible Detector monitoring at 311 nm wavelength, on a Phenomenex Luna Omega 5 µM Polar C18 250 x 21.2 mm column using gradient elution with using $H_2O$/MeOH with 0.1% TFA at 10 mL/min. The HPLC fractions of the major compound were concentrated under reduced pressure using a rotary evaporator, then lyophilized to dryness using a Labconco Freezone 4.5 L (- 84°C) lyophilizer and characterized by NMR which matched with the reported compound.[9] This product was analyzed for purity using RP analytical HPLC equipped with Waters 1525 with a 2489 UV/Visible Detector monitoring at 311 nm wavelength, on a Phenomenex Luna 5 µM C18(2) 250 x mm column using gradient elution with using $H_2O$/MeOH with 0.1% TFA at 1 mL/min. The major product was used for click chemistry.

Analytical HPLC chromatogram of 9-propargyl-2-amino-6-chloropurine

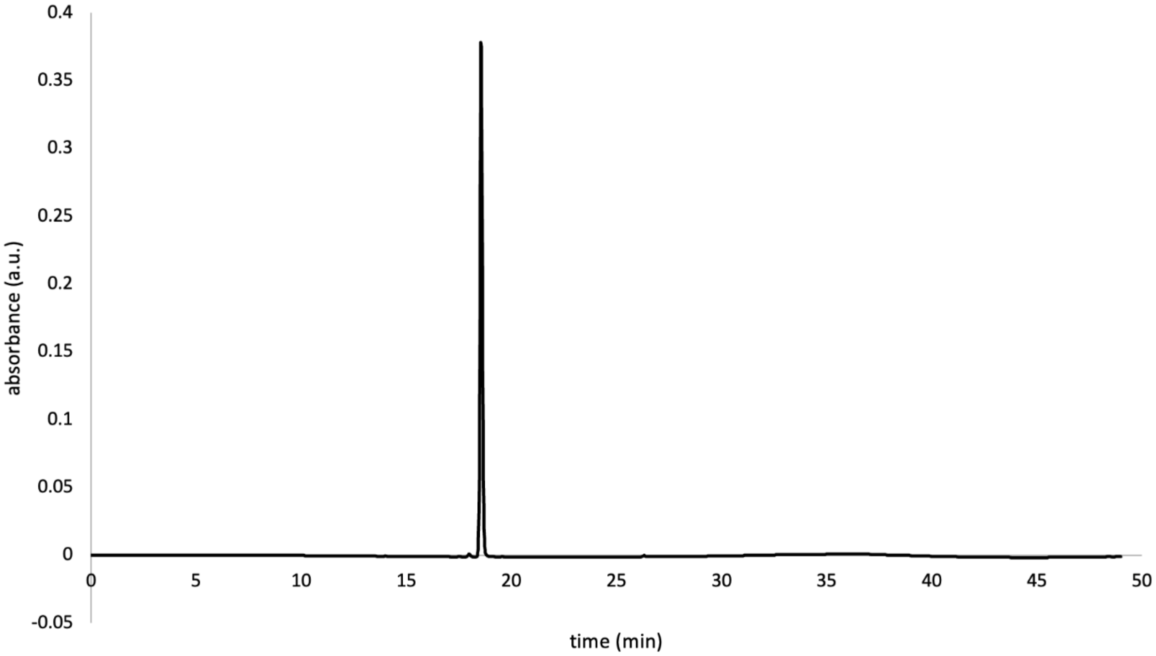

ESI High Resolution Mass Spectrum for 9-propargyl-2-amino-6-chloropurine

m/z calculated for $C_8H_6ClN_5$ $[M+H]^+$ 208.0385, found 208.0388.

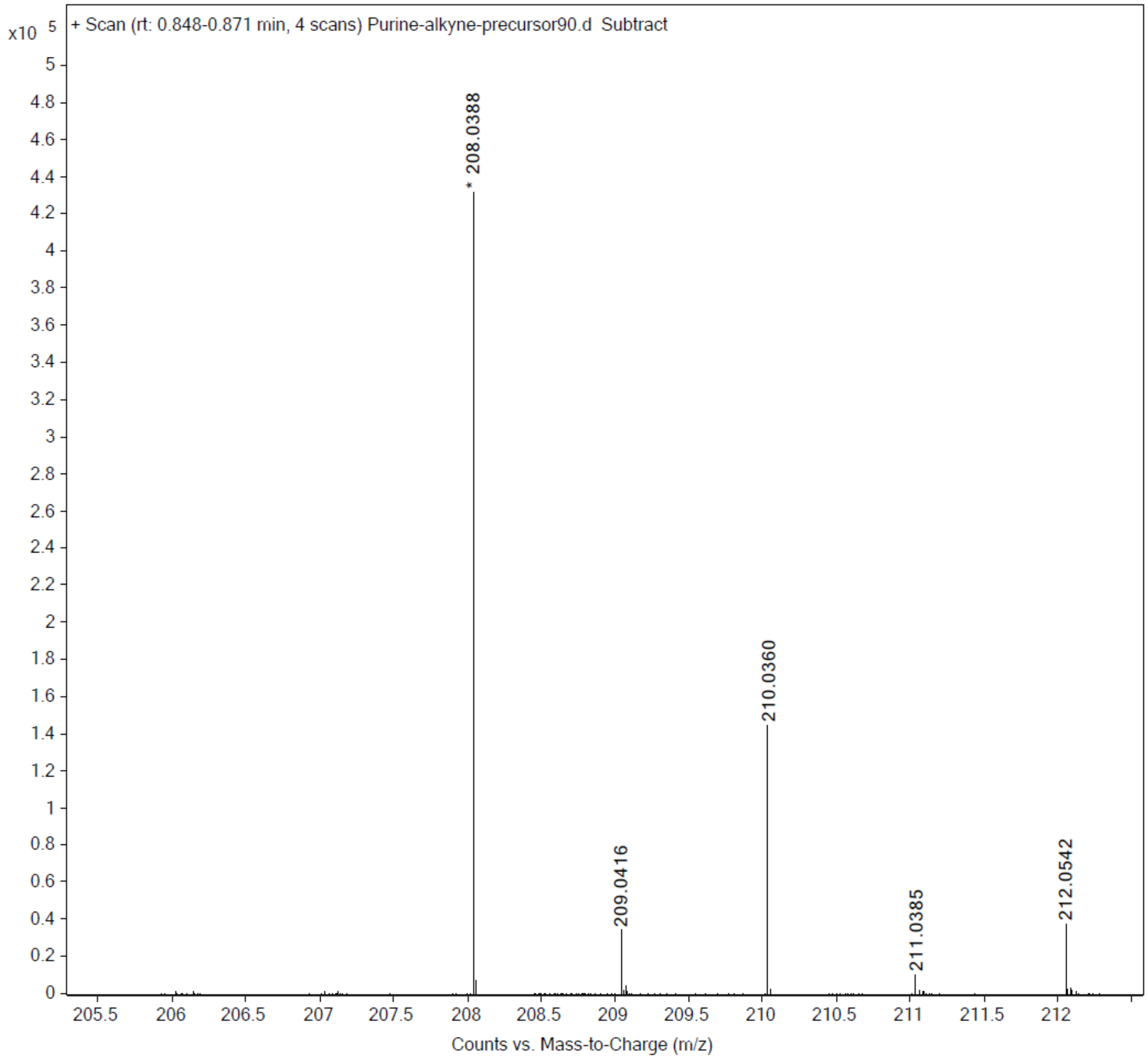


NMR of 9-propargyl-2-amino-6-chloropurine

**6-chloro-9-(prop-2-yn-1-yl)-9H-purin-2-amine**. White solid;[9] $^1$H NMR (600 MHz, DMSO-*d₆)* δ 8.18 (s, 1H, 8-H), 7.02 (brs, 2H, $-NH_2$), 4.93 (d, 2H, J=1 Hz, $-CH_2$), 3.48 (t, 1H, J = 1 Hz, C≡CH).

**Scheme S2. High-Throughput Synthesis of Triazole-Containing BIIB021 Derivatives**

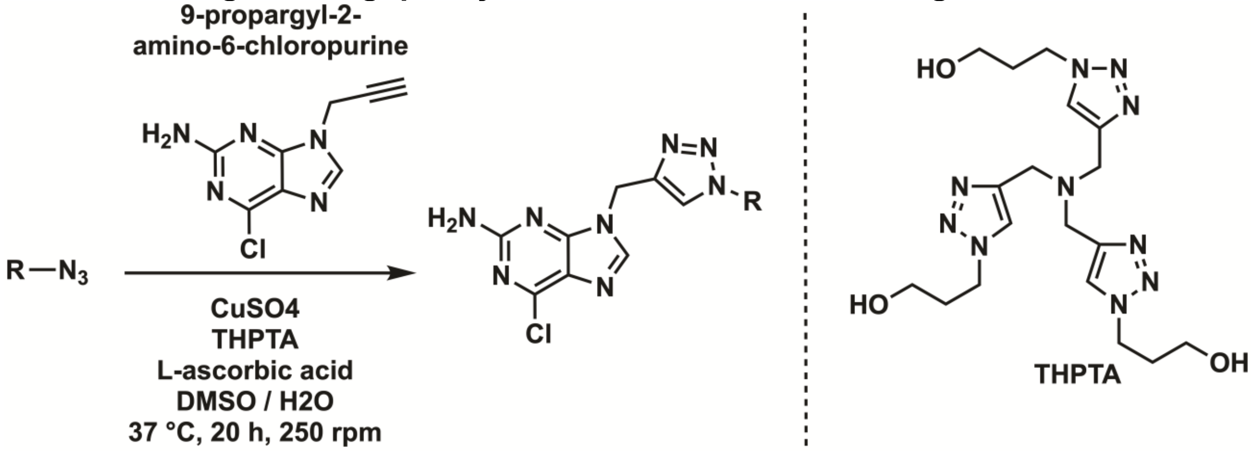


Triazole analogs were synthesized based on literature procedure.[10] Azide solutions from the azide library in Plates 1-5 were initially at a concentration of 100 mM in DMSO. Azides were added in each well of a 96-well plate at a concentration of 10 mM. To each well of this newly loaded plate, L-ascorbic acid solution was added to a concentration of 40 mM along with 10 mM of 9-propargyl-2-amino-6-chloropurine (synthesis shown in *Scheme S1*) and 2 mM of $CuSO_4$/THPTA in a solution of DMSO and water at a 3:2 ratio to a total volume of 100 µL. The plates were sealed and swirled at 250 rpm and 37°C for 20 hours to afford the corresponding triazole product in each well.

**Scheme S3. Synthesis of CliMB-325.**

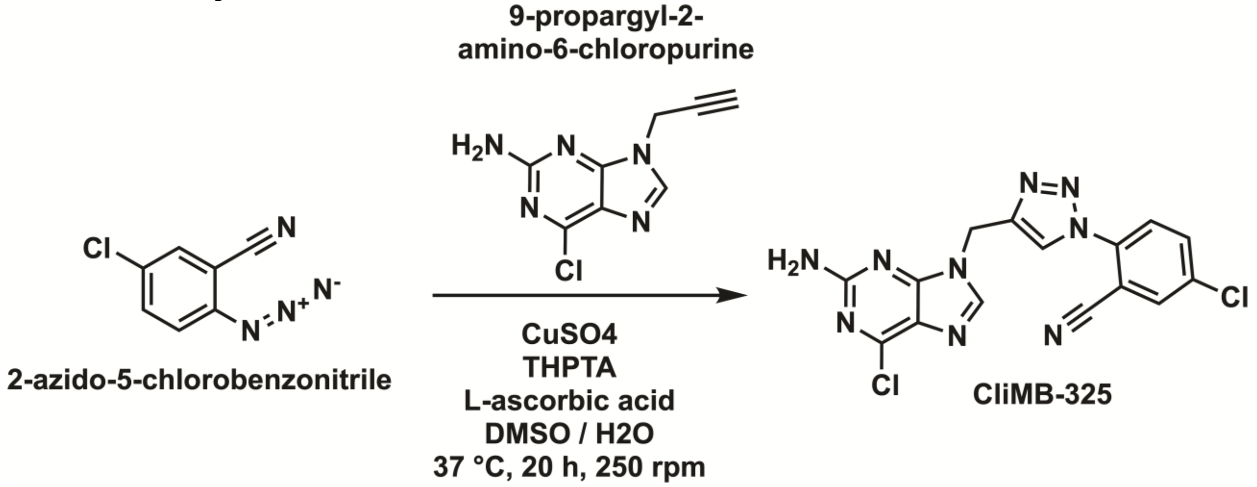


In a 50 mL conical tube, the following reagents were added: 10 mM of 2-azido-5-chlorobenzonitrile (azide #325 from 380 compound screen), 40 mM of aqueous L-ascorbic acid, 10 mM of 9-propargyl-2-amino-6-chloropurine (synthesis shown in *Scheme S1*), 2 mM of aqueous $CuSO_4$/THPTA solution, in a 3:2 ratio of DMSO to water at a total volume of 15 mL. The tube was swirled at 250 rpm and 37°C for 20 hours to yield **CliMB-325**. The compound was purified by reverse-phase preparative high-performance liquid chromatography (RP-HPLC) equipped with Waters 1525 with a 2489 UV/Visible Detector monitoring at 311 nm wavelength, on a Phenomenex Luna Omega 5 μM Polar C18 250 x 21.2 mm column using gradient elution with using $H_2O$/MeCN with 0.1% TFA at 10 mL/min. The HPLC fractions of the desired purified product were concentrated under reduced pressure using a rotary evaporator, then lyophilized to dryness using a Labconco Freezone 4.5 L (- 84°C) lyophilizer. The product was analyzed for purity using RP analytical HPLC equipped with Waters 1525 with a 2489 UV/Visible Detector monitoring at 311 nm wavelength, on a Phenomenex Luna 5 μM C18(2) 250 x mm column using gradient elution with using $H_2O$/MeCN with 0.1% TFA at 1 mL/min. The final product was stored at -20°C until further use, and stocks were made at 10 mM in DMSO.

Analytical HPLC Chromatogram of **CliMB-325**

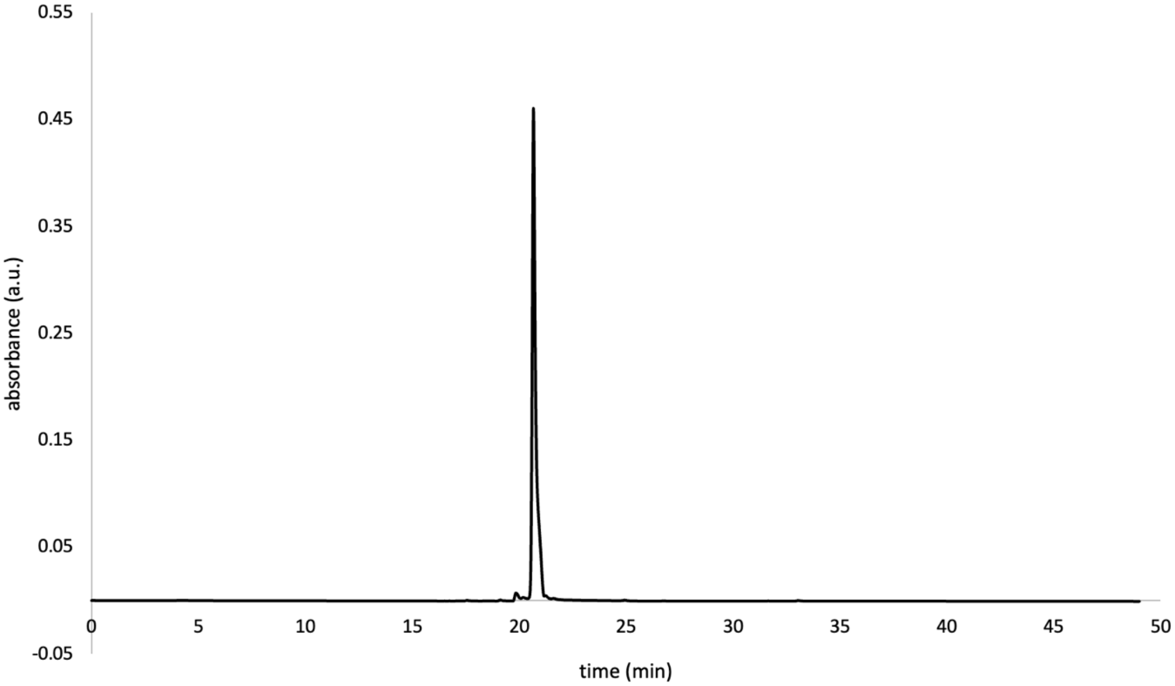

ESI High Resolution Mass Spectrum for **CliMB-325**

m/z calculated for $C_{15}H_{10}Cl_2N_9$ $[M+H]^+$ 386.0431, found 386.0439.

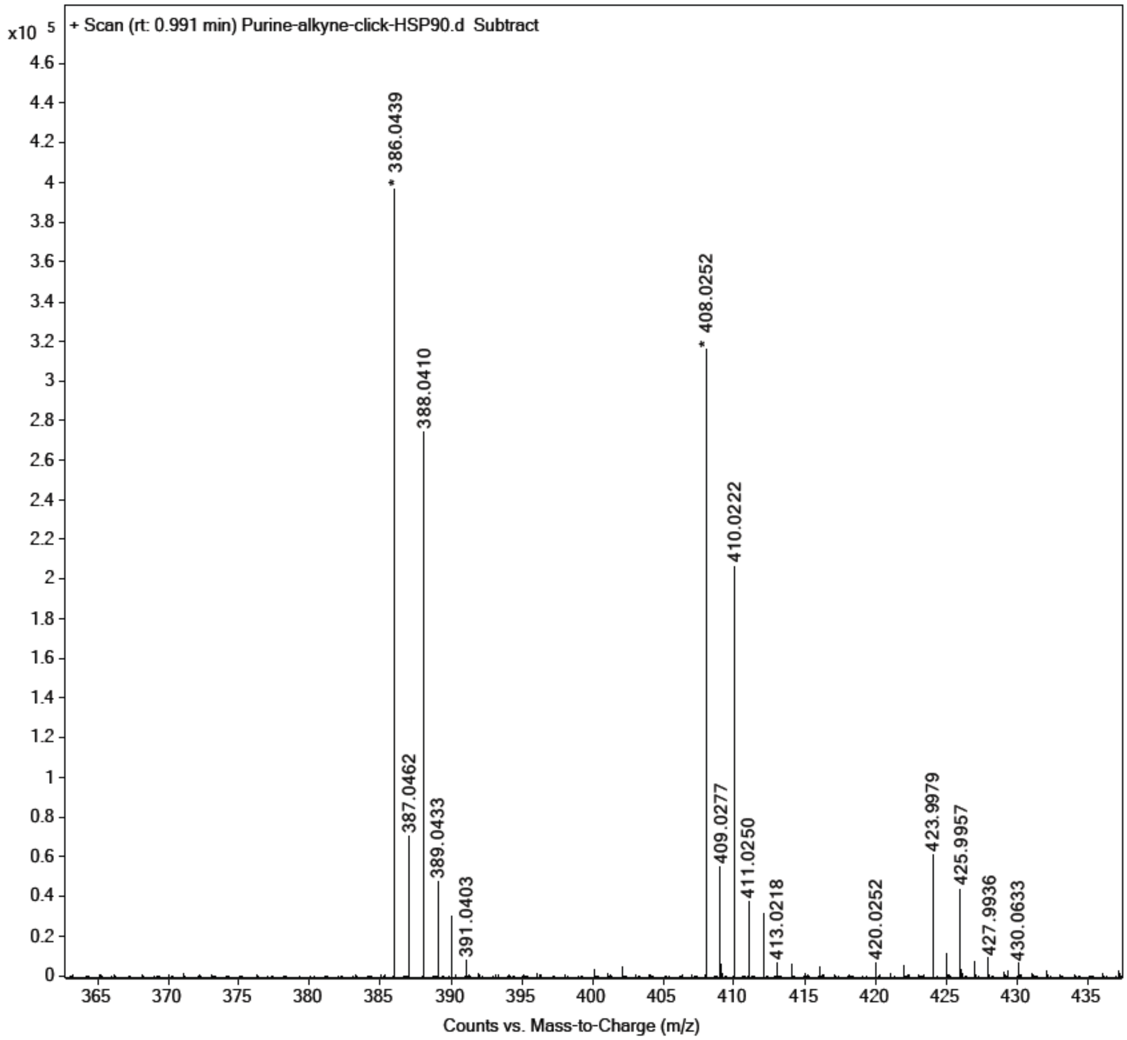

$^1$H NMR of **CliMB-325**

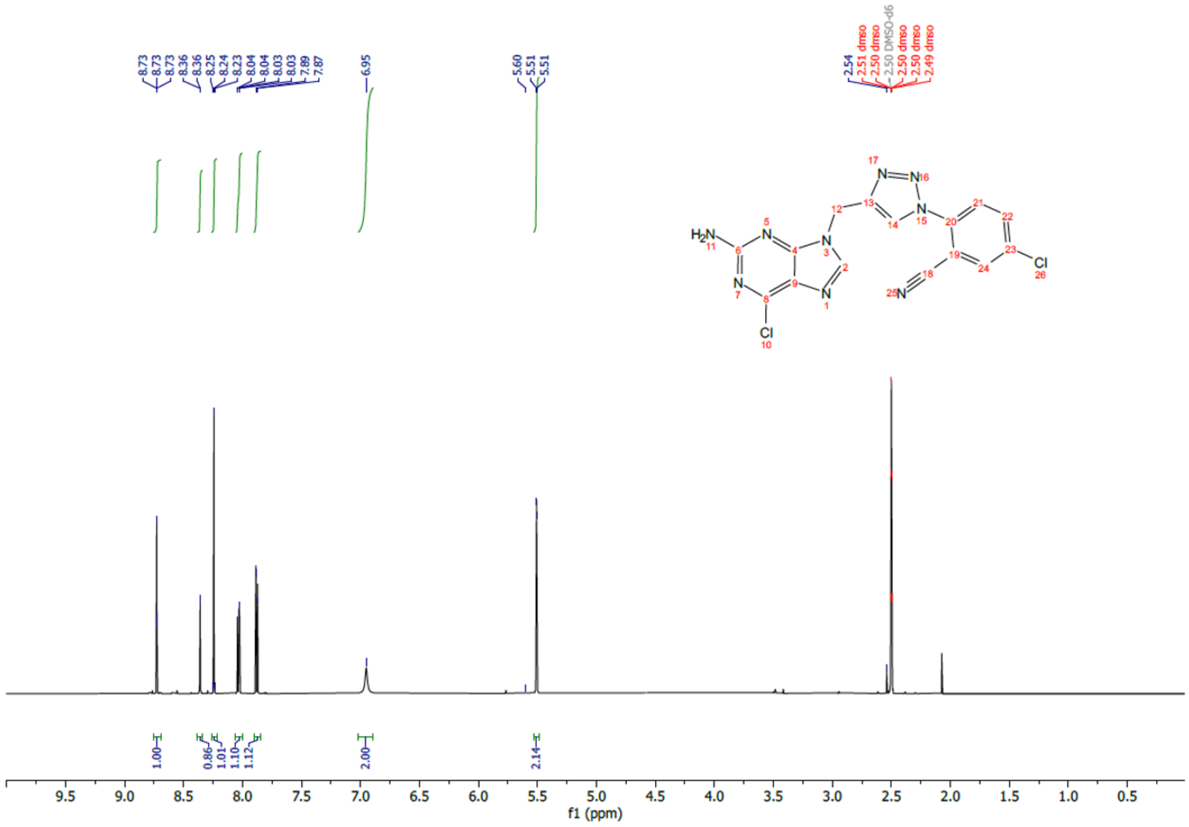

$^{13}$C NMR of **CliMB-325**

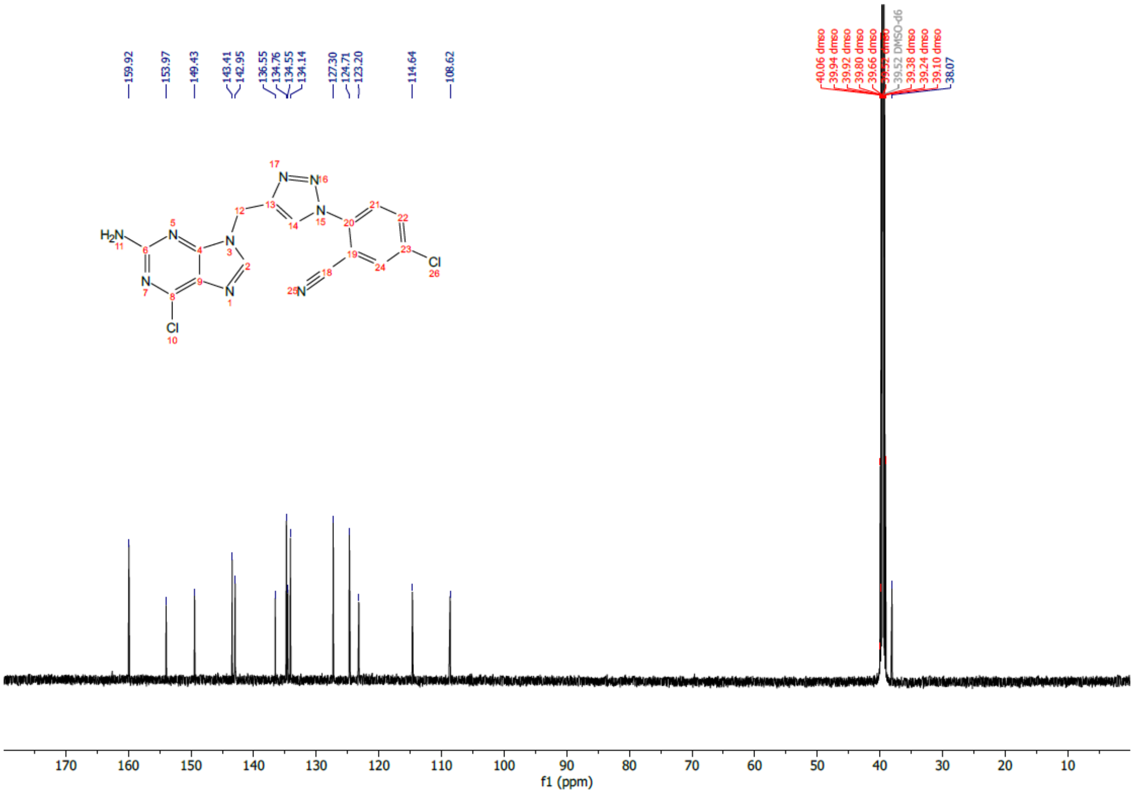


**2-(4-((2-amino-6-chloro-9H-purin-9-yl)methyl)-1H-1,2,3-triazol-1-yl)-5-chlorobenzonitrile**. White solid; $^{1}$H NMR (600 MHz, DMSO-*d₆)* δ 8.73 (s, 1H, 8-H), 8.36 (d, 1H, J= 1 Hz, Ar-H), 8.24 (s, 1H, triazole-H), 8.04 (dd, 1H, J= 3.6 Hz, Ar-H), 7.88 (d, 1H, J= 3.6 Hz, Ar-H), 6.95 (brs, 2H, -NH), 5.51 (s, 2H, $-CH_2$); $^{13}$C NMR (150 MHz, DMSO-$d_6$) δ 159.9, 153.9, 149.4, 143.4, 142.9, 136.5, 134.7, 134.5, 134.1, 127.3, 124.7, 123.2, 114. 6, 108.6, 38.0
